# Multi-Omic Analysis Reveals Lipid Dysregulation Associated with Mitochondrial Dysfunction in Parkinson’s Disease Brain

**DOI:** 10.1101/2024.07.18.604051

**Authors:** Jenny Hällqvist, Christina E Toomey, Rui Pinto, Anna Wernick, Mesfer Al Sharhani, Simon Heales, Simon Eaton, Kevin Mills, Sonia Gandhi, Wendy E Heywood

**Affiliations:** Translational Mass Spectrometry Research Group, Genetic & Genomic Medicine, UCL Great Ormond Street Institute of Child Health, WC1N 1EH London, United Kingdom; Queen Square Brain Bank for Neurological Disorders, UCL Queen Square Institute of Neurology, WC1N 1PJ London, United Kingdom; Department of Clinical and Movement Neurosciences, UCL Queen Square Institute of Neurology, WC1N 3BG London, United Kingdom; The Francis Crick Institute, NW1 1AT London, United Kingdom; MRC Centre for Environment and Health, School of Public Health, Department of Epidemiology and Biostatistics, Faculty of Medicine, Imperial College London, London W12 0BZ, United Kingdom; UK Dementia Research Institute at Imperial College London, London W12 0BZ, United Kingdom; MRC-NIHR BRC National Phenome Centre, Section of Bioanalytical Chemistry, Division of Systems Medicine, Department of Metabolism, Digestion and Reproduction, Faculty of Medicine, Imperial College London, Hammersmith Hospital Campus, London W12 0NN, UK; Neurometabolic Unit, National Hospital for Neurology and Neurosurgery, Queen Square & UCL Great Ormond Street Institute of Child Health, WC1N 3BG London, United Kingdom; College of Applied Medical Sciences, King Khalid University, Abha, Saudi Arabia; Developmental Biology and Cancer University College London Great Ormond Street Institute of Child Health, WC1N 1EH London, United Kingdom

**Keywords:** Parkinson’s Disease, brain tissue, phospholipids, sphingolipids, mass spectrometry, proteomics, multi-omics, mitophagy

## Abstract

Parkinson’s Disease (PD) is an increasingly prevalent condition within the aging population. PD can be attributed to rare genetic mutations, but most cases are sporadic where the gene-environment interactions are unknown/likely contributory. Age related dysregulation of the glycosphingolipid degradation pathway has been implicated in the development of PD, however, our understanding of how brain lipids vary across different regions of the brain, with age and in disease stages, remains limited.

In this study we profiled several phospho- and sphingolipid classes in eight distinct regions of the human brain and investigated the association of lipids with a spatio-temporal pathology gradient, utilising PD samples from early, mid, and late stages of the disease. We performed high-precision tissue sampling in conjunction with targeted LC-MS/MS and applied this to post-mortem samples from PD and control subjects. The lipids were analysed for correlations with untargeted proteomics and mitochondrial activity data, in a multi-omics approach. We concluded that the different brain regions demonstrated their own distinct profiles and also found that several lipids were correlated with age. The strongest differences between PD and controls were identified in ganglioside, sphingomyelin and n-hexosylceramides. Sphingomyelin was also found to correlate with several proteins implicated in Parkinson’s disease pathways. Mitochondrial activity was correlated with the levels of several lipids in the putamen region. Finally, we identified a gradient corresponding to Braak’s disease spread across the brain regions, where the areas closer to the brainstem/substantia nigra showed alterations in PC, LPC and glycosphingolipids, while the cortical regions showed changes in glycosphingolipids, specifically gangliosides, HexCer and Hex2Cer.

**Graphical Abstract:** 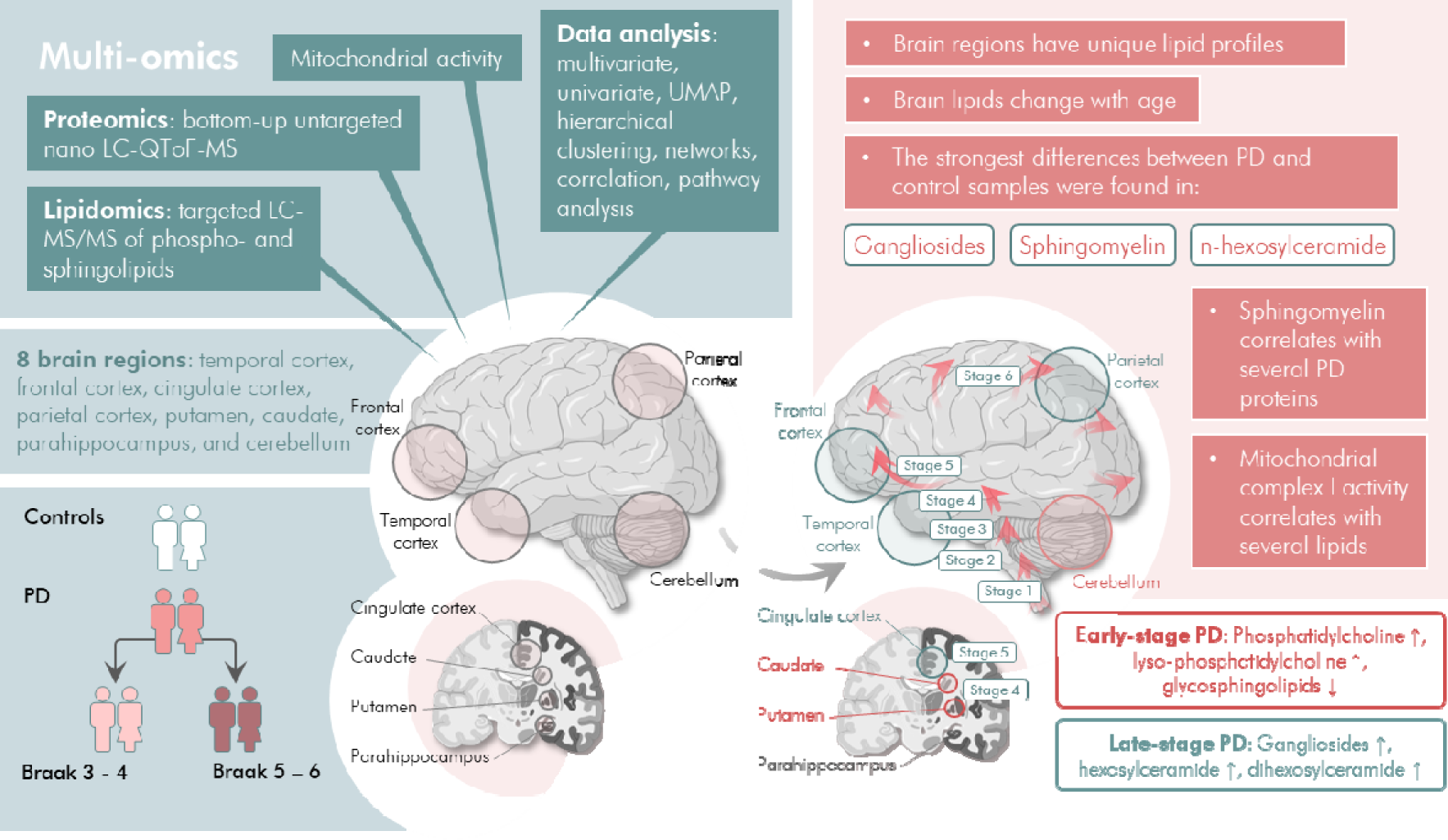

## Introduction

Parkinson’s Disease (PD) is becoming increasingly prevalent in our ageing population and is the most common neurodegenerative movement disorder, affecting 1–2% of people aged over 65, with an estimatedlJ>lJ6.1 million people affected worldwide ^1^.There has been a wealth of research into genetic and proteomic changes affected by PD^2–5^, however it is becoming more and more apparent that dysregulated lipid pathways also play a role in the aetiology of the disease^6–9^. Importantly, it has been observed that α-synuclein aggregation is affected by lipids^9^ and that α-synuclein inclusions have a high lipid content^10^. However, considering that the brain consists of 60 – 70% lipids, there is little recent literature^11,12^ to describe the normal human brain lipid profile, how it varies across different brain regions, age and disease state.

The relevance of lipid changes is more difficult to delineate than genes or proteins, and their functions are not yet completely understood. Lipids are highly heterogenous with many isoforms deriving from carbon chain length, functional groups, and the presence of unsaturated sites. The relevance and function of the many isoforms is yet to be understood, but in a broad sense, sphingolipids are highly enriched in the nervous system where they act as constituents of plasma membranes^13^ and play a role in membrane fluidity, recognition, signalling, and many cellular functions^14^. The major sphingolipid, sphingomyelin, has been implicated in PD^15^ particularly due to its role in α-synuclein exosome composition where it may be involved in mitigating α-synuclein spread ^16^. The glycosphingolipid degradation pathway is a key pathway implicated in genetic PD with defects in *GBA1* activity being the most common genetic cause for PD. A comprehensive figure of the glycosphingolipid degradation pathway is given in **Supplementary Figure S1**. Phospholipids belong to another class of lipids which are enriched in the brain and integral to the formation of plasma membranes. Dysregulation of phospholipids has been implicated in Alzheimer’s disease^17^ as well as following brain injury^18^, but they are also known to interact with α-synuclein^9^.

To expand on the knowledge of lipid expression in the human brain, we have profiled multiple brain regions in sporadic early-to mid-stage PD (Braak stage 3-4), and late-stage PD (Braak stage 5-6), exploring how the lipid profile changes across eight brain regions in relation to age and disease, and correlated these findings with proteomics data in a multi-omic approach. We applied targeted lipid panels consisting of the main non-cholesterol lipids that are abundant in the brain, including sphingolipids, glycosphingolipids, and phospholipids as well as their deacylated lyso-forms. We also included phosphatidylethanolamine plasmalogens, an ether glycerophospholipid derived from the peroxisome which is abundant in brain tissue and has been found to be reduced in lipid rafts in the frontal cortex of PD affected brains^19^. **Table 1** lists the lipids and isoforms included in this analysis.

**Table 1.** Compound classes measured and corresponding internal standards used. The table shows the analyte classes, abbreviations and monitored isoforms, and the internal standards used to normalise the abundancies. d: deuterium-labelled, LPC: Lyso-PC, LPE: Lyso-PE, LPE(P): Lyso-PE(P), LSM: Lyso-SM.

| Analyte compounds | Abbreviation | Isoforms | Internal standard |
| --- | --- | --- | --- |
| Ceramide | Cer d18:1 | 16-24 (+OH 16-24) | d<sub>3</sub>-ceramide d18:1/18:0 |
| Ganglioside GM1 | GM1 d18:1 | 16-24 | d<sub>3</sub>-GM1 d18:1/16:0 |
| Ganglioside GM2 | GM2 d18:1 | 16-20 | d<sub>3</sub>-GM2 d18:1/16:0 |
| Ganglioside GM3 | GM3 d18:1 | 18 | d<sub>3</sub>-GM3 d18:1/16:0 |
| N-acetylneuraminic acid | NANA | n/a | d<sub>3</sub>-GM1 d18:1/16:0 |
| Hexosylceramide (glucosylceramide + galactosylceramide) | HexCer d18:1 | 16-26 (+OH 18-26) | d<sub>3</sub>-glucosylceramide d18:1/16:0 |
| Di-hexosylceramide (lactosylceramide + galabiosylceramide) | Hex2Cer d18:1 | 16-24 (+OH 24) | d<sub>3</sub>-glucosylceramide d18:1/16:0 |
| Globotriaosylceramide | Gb3Cer d18:1 | 16-18 | d<sub>3</sub>-glucosylceramide d18:1/16:0 |
| Globoside | Gb4Cer d18:1 | 18 | d<sub>3</sub>-glucosylceramide d18:1/16:0 |
| Globotriaosylsphingosine/lyso-Gb3 | Gb3Sph | n/a | N-glycinated globotriaosyl-sphingosine d18:1 |
| Phosphatidylcholine | PC | 34-40 | N-glycinated globotriaosyl-sphingosine d18:1 |
| Phosphatidylethanolamine | PE | 34-40 | N-glycinated globotriaosyl- sphingosine d18:1 |
| PE plasmalogen | PE(P) 16:0, 18:0, 18:1 | 18-22 | N-glycinated globotriaosyl-sphingosine d18:1 |
| Lyso-phosphatidylethanolamine | LPC | 16-22 | N-glycinated globotriaosyl-sphingosine d18:1 |
| Lyso-phosphatidylethanolamine | LPE | 16-22 | N-glycinated globotriaosyl-sphingosine d18:1 |
| Lyso-phosphatidylethanolamine PE | LPE(P) | 16-18 | N-glycinated globotriaosyl-sphingosine d18:1 |
| Sphingomyelin | SM | 34-36 | N-glycinated globotriaosyl-sphingosine d18:1 |
| Lyso-sphingomyelin | Lyso-SM | 16-18 | N-glycinated globotriaosyl-sphingosine d18:1 |
| Lyso-sphingomyelin 509/ N-palmitoyl-O-phosphocholineserine | Lyso-SM (509) | n/a | N-glycinated globotriaosyl-sphingosine d18:1 |
| Hexosylsphingosine /Lyso-Gb1 (glucosylsphingosine + galactosylsphingosine) | HexSph | n/a | d<sub>5</sub>-glucosylsphingosine d18:1 |
| Dihexosylsphingosine/Lyso-Gb2 (lactosylsphingosine galabiosylsphingosine) | Hex2Sph | n/a | d<sub>5</sub>-glucosylsphingosine d18:1 |

## Results

### Lipid expression differs between anatomical regions and is modulated by age in healthy brains

We first examined the group of control samples (n = 107, 38% F) to establish a baseline expression of the analysed lipids in healthy, aged brains, not affected by PD. The average distribution of each lipid class showed that there were spatial differences in which respective lipids expressed more strongly. Examining the lipid classes in detail, we found that ceramide was the lipid class changing the least between regions, with a variation of 15%, while lyso-phosphatidylethanolamine (lyso-PE) and n-hexosylceramide (the sum of glucosylceramide and galactosylceramide) varied the most, presenting coefficients of variation of 82% and 52%, respectively (**Figure 1A**). We applied hierarchical clustering to evaluate the correlation between lipid-levels across the brain to determine whether the regions shared similar expression profiles (**Figure 1B**). Three regional clusters formed, consisting of (i) frontal cortex (FrC), caudate (CA) and parahippocampus (PHp), (ii) cerebellum (CBM), cingulate cortex (CiC) and putamen (PU), and (iii) parietal (PtC) and temporal cortices (TCtx). The primary driving parameter for this clustering was found to be a general elevated expression of both phospho- and sphingolipids in the FrC-CA-PHp cluster, while the CBM-CiC-PU cluster demonstrated elevated levels of ganglioside, especially prominent in CiC and PU. Lastly, the PtC-TCtx cluster was distinguished by lower levels of most lipids but lyso-phospholipids, and ceramide. The clustering also showed that sphingomyelin, n-hexosylceramide (where n represents the number of hexosyl groups) and *N*-acetylneuraminic acid presented similar expression profiles, as did the diacylated phospholipids. We further modelled the lipids in a Uniform Manifold Approximation and Projection (UMAP)^20^ to obtain an overview of how the different lipids and lipid classes related to each other. Five main clusters emerged, consisting of (i) gangliosides (GM1-3) and dihexosylceramide (Hex2Cer), (iii) phospholipids, (iv) lyso-phospholipids, (v) HexCer, and (vi) ceramide and plasmalogen PE (**Figure 1C**).

**Figure 1.**
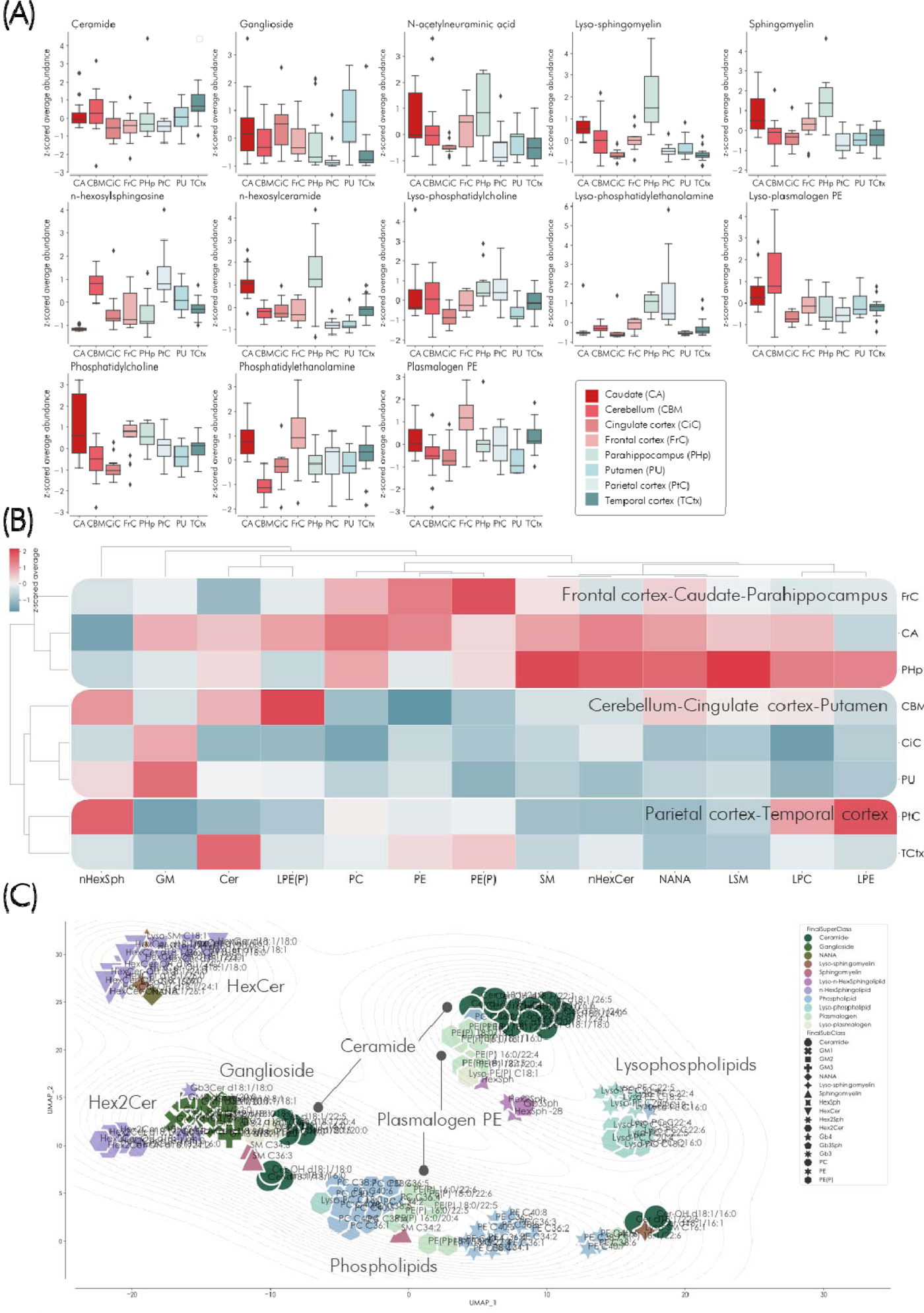
Lipid expression in control brains without brain pathology. (A) Box and whisker plots displaying the z-scored sum of lipids per class and the distribution in the different regions. The whiskers show the minimum and maximum and the boxes show the 25^th^ percentile, the median and the 75^th^ percentile. Values outside 1.5 times the interquartile are represented by dots. It was demonstrated that the phospholipids, ceramide and sphingomyelin displayed an even distribution between the different regions. (B) Hierarchical clustering showing the average of each compound per region in controls. Three main clusters formed consisting of (i) frontal cortex, caudate and parahippocampus, (ii) cerebellum, cingulate cortex and putamen, and (iii) parietal and temporal cortices. The primary driving parameter for this clustering was found to be an elevated expression of most lipids in the frontal cortex-caudate-parahippocampus cluster. The clustering method was set to average, with cosine as distance metric. (C) UMAP projection demonstrating how the lipid classes relate to each other. The projection grouped into five main clusters, largely made up of lipids from the same class although overlap between classes was also observed. The UMAP points were coloured according to main class and shaped according to sub-class. The projection was set to model 12 neighbours (average lipid class size) with Euclidean distance metric.

To determine whether age affects the brain lipid profile, we examined the control samples and noted that n-hexosylceramide was affected across all brain regions, apart from the cingulate and frontal cortices, and increased in abundance with increasing age (**Supplementary Results Figure S2**). We also evaluated sex-related variations in the lipid profiles by Orthogonal Partial Least Squares – Discriminant Analysis (OPLS-DA) but did not find any significant differences in expression (ANOVA CV p = 1).

In conclusion, we found that the lipid profile differs regionally across the brain in controls and that age influences the lipid expression. The caudate and the temporal cortex were the regions most affected by age and were found to be particularly susceptible to in age-related lipid changes, specifically in n-hexosylceramide and ceramide. These findings underscore the importance of comparing different regions when performing lipidomic analyses rather than using whole brain homogenate, as the regional lipidomes differ drastically.

### Age adjustment

To avoid interpreting differences in age between the groups as relevant to discriminating between PD and controls, given the significant correlations between age and several of the lipids in the control group, we adjusted the data region-wise for age before performing any further analyses. Only the variables significantly correlated with age were adjusted and evaluated in both a total- and a region-wise OPLS model with age set as the y-vector to verify that the adjustment was effective. None of these models were significant, thereby demonstrating that the age-adjustment was performed successfully. The p-values showing if a variable was significantly related to age are shown in **Supplementary Results Table S1**.

### Ceramide, sphingomyelin and ganglioside levels drive the changes observed in the PD brain lipid profile

We next explored differences between PD and control. We constructed a PCA model of the PD and control samples throughout the brain regions. The first and second principal components largely separated the samples into controls and PD and the corresponding loadings showed that gangliosides were elevated in PD, along with long-chained and poly-unsaturated phospholipids (**Figure 2A-B**). The third and fourth principal components described differences in brain regions and, in similarity to what we found in the control samples, the PCA loadings demonstrated that the eight different PD brain regions were enriched in different lipid classes. In likeness to what we had observed in the control samples alone, we noted that the cortices were abundant in ceramide, phospholipids and plasmalogen PE, while the inner brain regions were richer in ganglioside and lyso-n-hexosylsphingosine (including glucosylsphingosine and psychosine) (**Figure 2C-D**). To evaluate the overall lipid expression related to the PD and control groups, we adjusted the data for brain regions before modelling the classes by discriminant OPLS-DA (**Figure 2E-F**). This model was found to be highly significant (ANOVA CV p = 5.4 E^-12^) and no regional influence could be discerned. The predictive loadings showed that the lipids most strongly responsible for the difference between the control and PD groups were higher levels of the ceramides C16 and C18 and lower levels of ceramide C22 and C24 in the PD group. All the measured gangliosides GM1, GM2 and GM3 were elevated in PD, as were sphingomyelin and lyso-sphingomyelin. Lyso-PE was lower in PD while lyso-phosphatidylcholine (lyso-PC) was higher. Phospholipids and plasmalogen PE were generally elevated in PD. Notably, the C22:6 fatty acid (docosahexaenoic acid, DHA) was found enriched in the upregulated phospholipids and is of particular interest due to the protective and anti-inflammatory properties of this specific fatty acid moiety.

**Figure 2.**
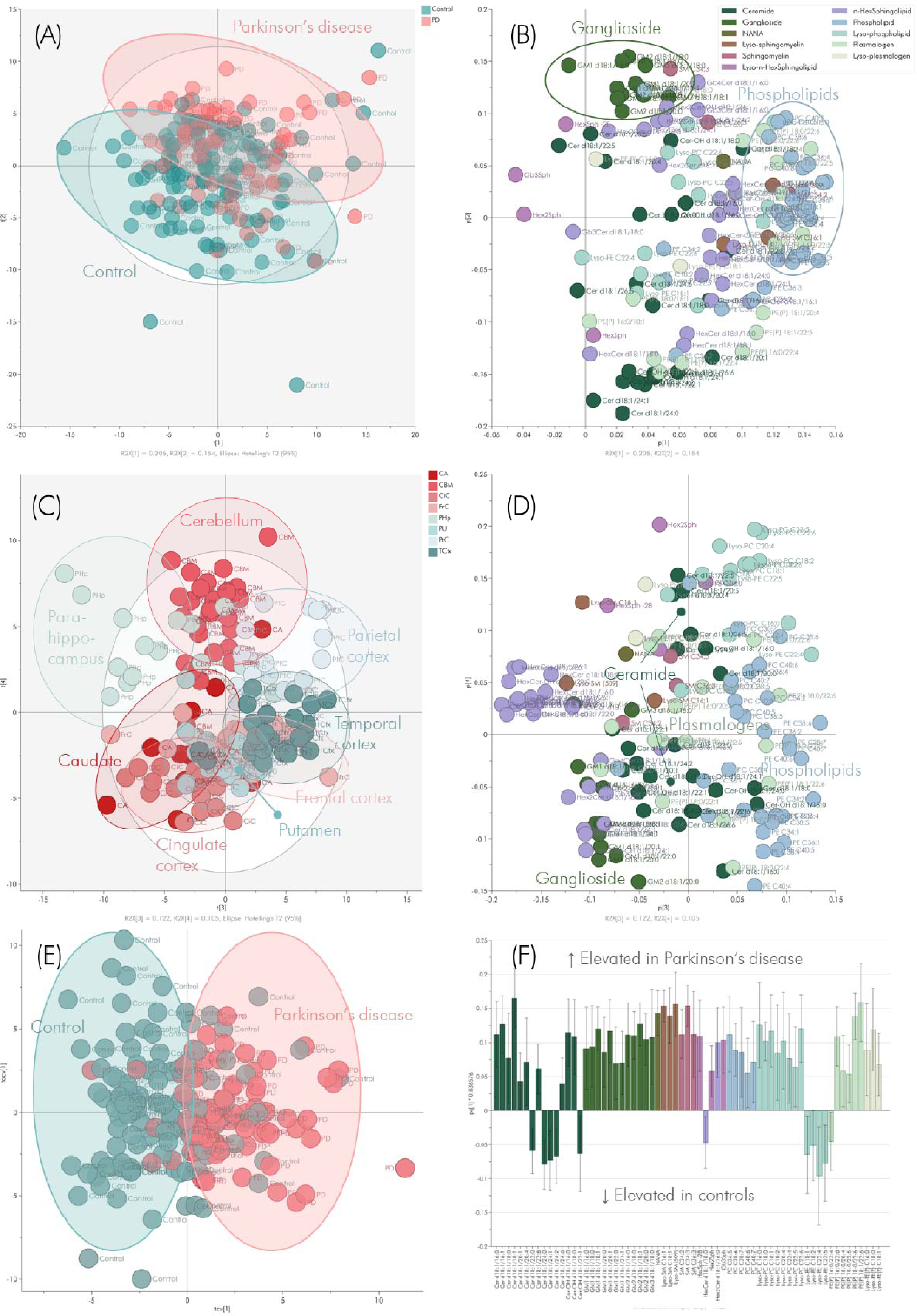
Multivariate analysis of PD and control. (A) PCA scores, the first (21% model variation) and second (15% model variation) components largely separated the samples into PD and control. (B) The corresponding loadings showed that gangliosides were elevated in PD, along with long-chained and poly-unsaturated phospholipids. (C) PCA scores from the third (12%) and fourth (11%) components grouped the samples into anatomical brain regions, where the putamen and cingulate cortex overlapped with caudate and the frontal cortex. The other regions generally formed well-defined clusters. (D) The corresponding loadings demonstrated that the eight different PD brain regions were enriched in different lipid classes. The cortices were rich in ceramide, phospholipid and plasmalogen, while the inner brain regions were richer in ganglioside and lyso-n-hexosylsphingosine. (E) The cross-validated scores from a region-adjusted OPLS-DA model of PD versus control, showing a clear separation between the groups. (F) Predictive OPLS-DA loadings showing the significant lipids where positive values represent lipids elevated in the PD group and negative values represent lower levels in PD. The model was significant with ANOVA CV p = 5.4 E^-12^ and permutations p < 0.001.

We further assessed the univariate and region-wise differences between PD and controls. We constructed signed (based on fold change direction) -log_10_ p-values and built a hierarchical cluster to summarise the differences (**Figure 3A**). The cingulate, temporal, frontal and parietal cortices formed one cluster, and the putamen, caudate and cerebellum another. This clustering seemingly showed a split across the anatomical regions with “Cluster 2” regions being closer to the substantia nigra, thereby conceivably following Braak’s suggested disease spread across the brain, whereas “Cluster 1” contained the brain regions furthest from the substantia nigra (**Figure 3B**). In Cluster 1, gangliosides and several HexCer and Hex2Cer isoforms were elevated in PD, while most phospholipids were lower. In Cluster 2, phospholipids, particularly PC and lyso-PC, were higher in PD than control, especially in the putamen and cerebellum, while glycosphingolipids were generally found lower. The parahippocampus did not group with either cluster but instead exhibited its own profile consisting of elevated levels of hydroxylated n-hexosylceramide, and low levels of most lyso-phospholipids and gangliosides.

**Figure 3.**
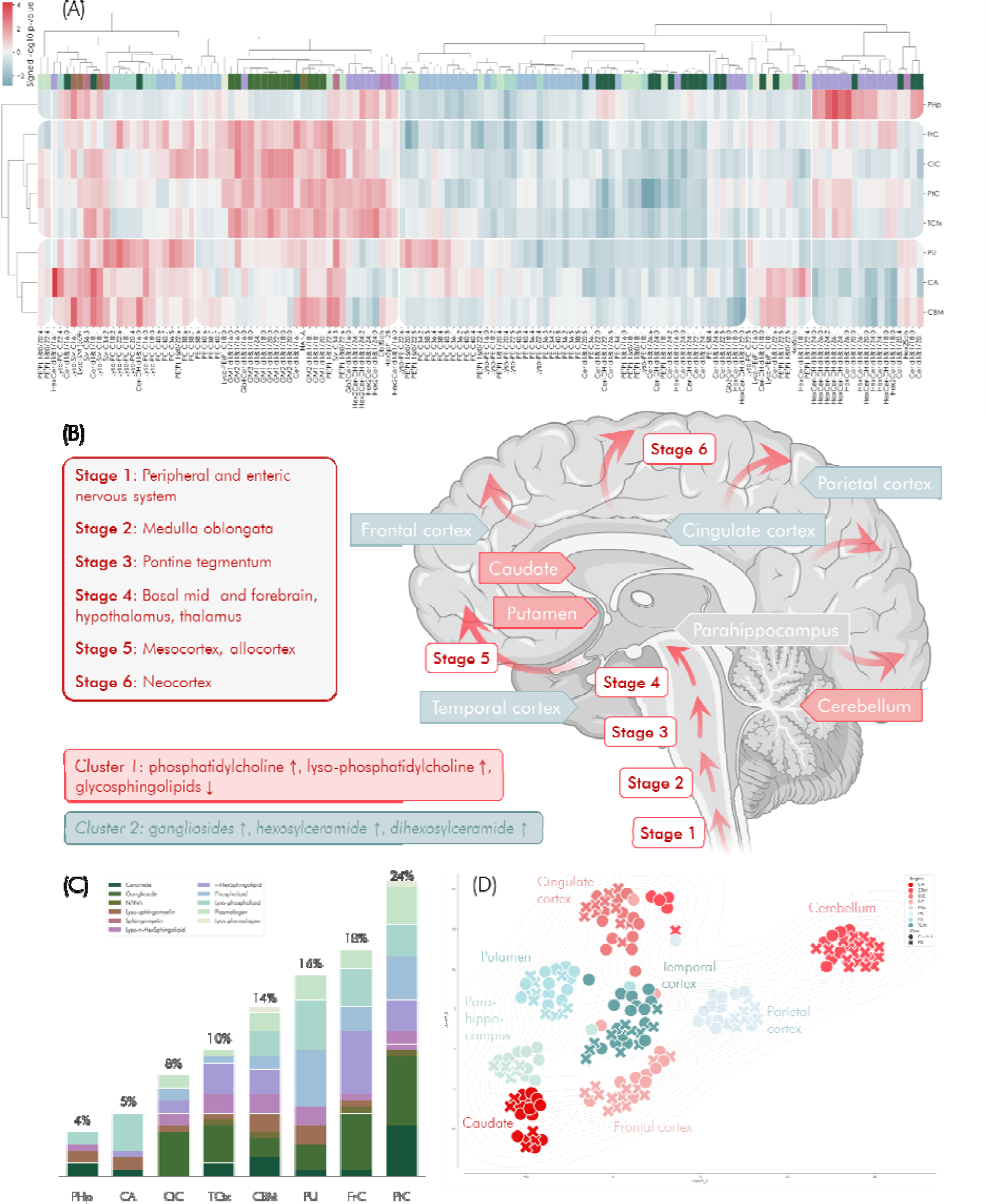
Differences in lipid brain expression between Parkinson’s disease and controls. (A) Hierarchical clustering of the expression differences (-log10 p-value) identified between control and PD in the eight different brain regions determined by Student’s t-test or Mann-Whitney’s U-test. Two main clusters formed consisting of (i) frontal, cingulate, parietal and temporal cortices, and (ii) putamen, caudate and cerebellum while the parahippocampus did not group with either cluster. The clustering method was set to average, and the metric was cosine. Red represents an increase, and blue a decrease, in PD. (B) The spread of pathology throughout the brain in Parkinson’s disease as described by Braak, and the changes identified in this study. We found two main clusters, the first cluster consisted of the more peripheral temporal, frontal, parietal cortices and the cingulate cortex, where we identified elevated levels of gangliosides, hexosylceramide and dihexosylceramide. The second cluster consisted of elevated levels of phospholipids and decreased levels of glycosphingolipids in the regions caudate, putamen and cerebellum. (C) Distribution of differentially expressed lipids in the brain regions, labelled by the total percentages of the nominally different lipids. The region with most numerous changes was the parietal cortex (24% of the total changes), followed by the frontal cortex (18%) and the putamen (16.0%). Ganglioside and n-hexosylated ceramide demonstrated the most abundant changes, while ceramide and lyso-SM were found altered in six out of seven regions. Phospholipids demonstrated the most numerous changes in the putamen and the parietal cortex and sphingolipids in the frontal and parietal cortices. (D) UMAP projection showing how the brain regions relate to each other. The regions generally formed distinct clusters separated from each other. Especially the cerebellum was distinctly different from the other regions. The difference between the regions was found to be greater than the difference between PD and control samples. The UMAP was set to model 23 neighbours (average region sample size), and the metric was Euclidean.

We evaluated the total number of changed compounds throughout the different regions in the whole dataset and noted that the parietal cortex was the region with the largest number of altered lipids between PD and control (24% of the total lipids), followed by the frontal cortex (18%) and the putamen (16%). When assessing the compound classes for changes between PD and control, we found that ceramide and lyso-SM were the most consistently changed lipids, both altered in seven out of eight regions of the brain (ceramide was unaffected in cingulate cortex and lyso-SM was unaffected in the parietal cortex). Lyso-PC and the glycosphingolipid classes n-hexosylceramide and ganglioside were also found to exhibit differences in most of the regions (**Figure 3C**).

We next used the signed -log_10_ p-values to project the samples in a UMAP. We found that samples from the same region generally clustered together and that the inner regions caudate, putamen and parahippocampus were located on the left side, the cingulate, temporal, parietal and frontal cortical regions were in the middle of the projection, while the cerebellum was separated to the right from the other regions. The regional clustering emphasises that the regions have different lipid profiles, but also that regions located spatially near each other also share similarities. No distinct separation could be identified between PD and control samples signifying that the regional differences are greater than the differences between healthy brains and PD (**Figure 3D**).

In summary, the parietal cortex, frontal cortex and the putamen were the regions showing the largest differences between PD and healthy controls. We found that ceramide, n-hexosylceramide, ganglioside, lyso -SM and lyso-PC were altered in the greatest number of regions

### The Braak stage lipid expression suggests a disease progression gradient

The samples had been categorized by Braak staging, classifying them as early-(Braak 3 - 4) or late-stage PD (Braak 5 - 6). Differences between PD groups and controls were assessed in three regions which had sufficient sample numbers in both early- and late-stage PD. These regions were cerebellum, frontal cortex, and putamen (details in **Supplementary Methods Table S1**). According to Braak’s map of pathology-spread through the brain at different disease stages, the putamen would be region most affected by pathology at early-stage PD, whereas the frontal cortex would not demonstrate pathology at this stage although it would in late-stage PD, and the cerebellum would not display any pathological aggregation. In the comparison of early-stage PD and controls, we could only identify a limited number of significant differences, consisting of higher ceramide and lyso-sphingomyelin levels in the cerebellum in PD, and higher levels of plasmalogen PE and lyso-PC in PD frontal cortex. Apart from these differences, the early PD group was statistically indistinguishable from controls. The putamen was the region which displayed the greatest number of significant differences in late-stage PD, with both lyso and non-lyso sphingo- and phospholipids altered. The frontal cortex also showed several changes in late-stage PD and, similarly to the putamen, these consisted of increases in both phospho- and sphingolipids (**Figure 4**). In addition, the putamen also demonstrated a large increase in sphingomyelin and lyso-sphingomyelin which was not seen in the frontal cortex. Further differences between the frontal cortex and the putamen were increased HexCer and lower levels of lyso-PE in the frontal cortex. Even though we were only able to distinguish small changes between early-stage PD and controls on a statistically significant level, we observed a step-like behaviour in most of the compounds where the early-stage PD samples were intensity-wise positioned in the middle between the control and late-stage PD groups. It is probable that the lack of significance in the early-stage PD group is due to the limited sample size, and consequently insufficient statistical power.

**Figure 4.**
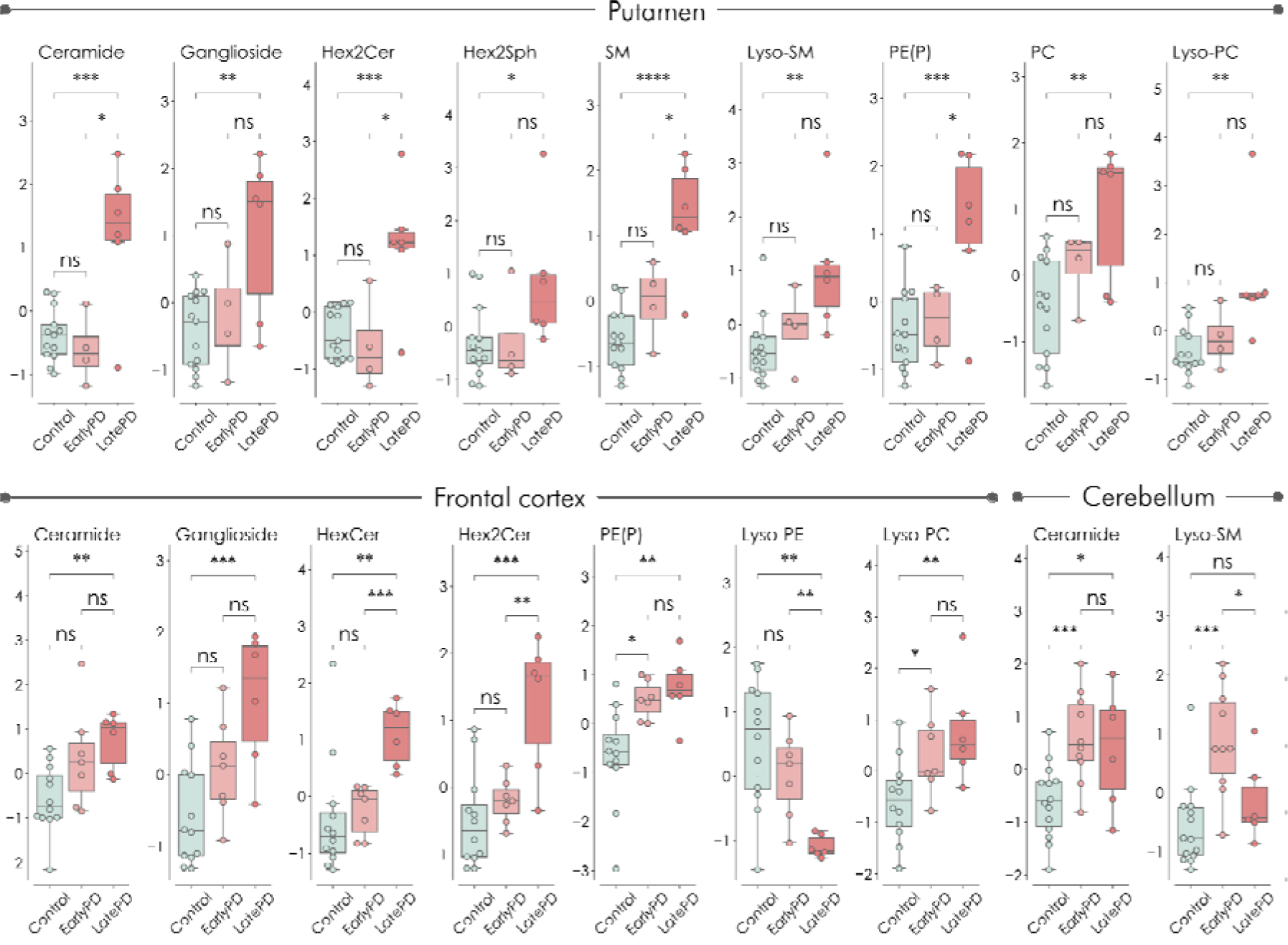
Box and swarm plots of the lipid expression in control, early-stage PD with Braak stages 3 – 4, and late-stage PD with Braak stages 5 – 6. The average of the statistically significant compounds post Benjamini-Hochberg multiple testing correction (FDR = 10%) are shown. The expression is displayed for each of the regions cerebellum, frontal cortex, and putamen. The whiskers show the minimum and maximum and the boxes represent the 25^th^ and the 75^th^ percentile, and the median. Values outside 1.5 times the interquartile are represented by dots. * p < 0.05, ** p < 0.01, *** p < 0.001, **** p < 0.0001, ns = non-significant.

In summary, the only region where we distinguished significant differences between control and early PD was the cerebellum, the differences consisting of elevated levels of ceramide and lyso-SM in PD. In the frontal cortex and the putamen, we observed an “intensity-ladder”, going from control levels to slightly elevated levels in early PD, to strongly elevated levels in late PD – thereby indicating that the lipids change with disease progression.

### Multi-omic analysis correlates sphingomyelin-levels in the putamen with proteins altered in PD

We next evaluated the correlation between the protein expression in the brain samples and the measured lipids, using Spearman correlation. Untargeted proteomics was used to profile these same samples, detailed in the publication by Toomey et al^21^. Significant correlations passing a

Benjamini-Hochberg FDR threshold set to 5% and proteins correlating with at least two lipids from the same compound class were considered. The results showed that three sphingomyelin isoforms significantly correlated with the expression of several proteins, positively with 14 proteins and negatively with 29 proteins, while in total, 25 of these proteins were nominally significant between PD and control in the proteomic data alone. We extracted the KEGG pathways associated with the lipid-correlated proteins and found that “Parkinson’s Disease” was the strongest enriched pathway followed by several other pathways of neurodegenerative disorders (**Figure 5**). Gene ontology (GO) suggested that biological processes related to mitochondrial energy metabolism were enriched.

**Figure 5.**
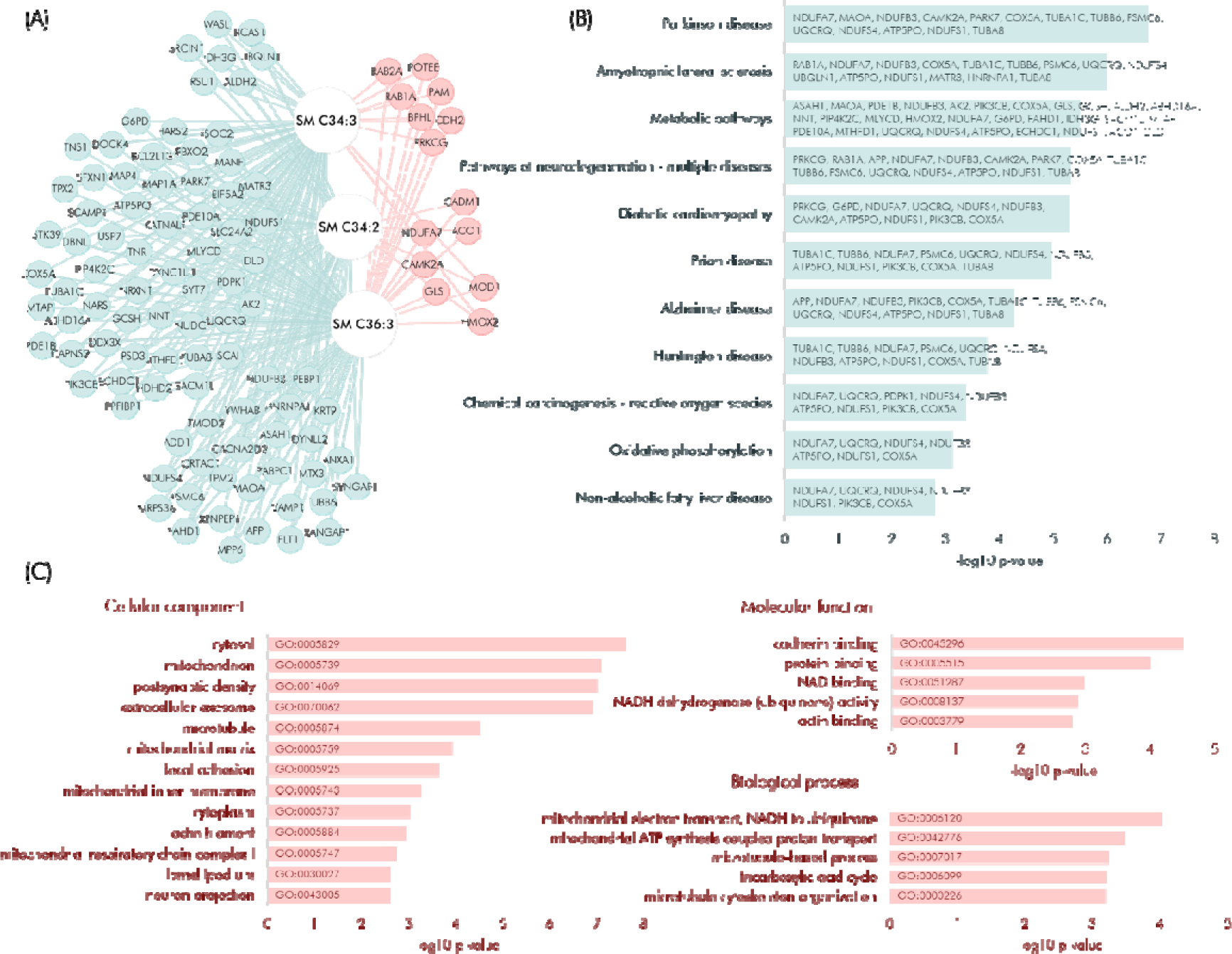
Correlation between lipids and proteins. The relationships between lipids and proteins in the regions putamen, parahippocampus and frontal cortex were evaluated by Spearman correlation, applying the Benjamini-Hochberg multiple testing correction procedure with FDR at 5%. The correlations were restricted to the ones significant in respective region post multiple testing correction, and only proteins which correlated with at least two lipids from the same compound class were considered. Only sphingomyelin in the putamen demonstrated significant differences between PD and controls, and significant lipid-protein correlations. (A) The network shows the significant interactions identified between proteins and sphingomyelin species in the putamen. Blue nodes represent negative correlations and red nodes positive correlations. Enriched pathways and GO annotations were extracted for significantly correlated proteins. The significantly enriched KEGG pathways (passing Benjamini-Hochberg FDR = 5%) are shown in (B) and demonstrate Parkinson’s disease as the top hit together with several other neurodegenerative conditions. (C) shows the significantly enriched GO cellular component terms, and the top five GO molecular function and biological process terms.

The correlation of sphingomyelin with several PD-implicated proteins suggests that there exist disease-specific interactions between lipids and proteins and highlights the importance of bringing together multiple omics platforms to identify more specific correlations and changes related to disease.

### Lipids in the putamen show significant correlations with mitochondrial complex I activity

We wanted to explore if any of the lipids were significantly correlated to markers of mitochondrial activity, given that the lipid-protein correlation-based pathway analysis suggested that mitochondrial energy activity was enriched. The mitochondrial complexes I, II-III, IV and citrate synthase were measured in the regions frontal cortex, parietal cortex and putamen as previously described^21^ and it was found that all of the complex ratios I, II-III, and IV to citrate synthase were significantly different between the PD and control groups in the putamen region, with complex-ratios I and II-III lower in PD, and complex-ratio IV higher in PD (**Supplementary Results Figure S3**). The relationship between the lipids and the ratios of the mitochondrial complexes to citrate synthase were evaluated in a linear mixed effects model where the brain regions were modelled as a random effect. The p-values from the interactions between the class variable (PD or control) and the complex proteins ratioed to citrate synthase were adjusted for multiple testing by the Benjamini-Hochberg procedure, and statistical significance accepted up to FDR = 10% (**Supplementary Results Table S2**). The results were visualised in a network (**Figure 6**) where we found that several lipids, consisting of a large proportion of the compounds in the plasmalogen PE and lyso-PC classes, and also of ceramide, HexCer and HexSph, were significantly different between PD and control in the regression of complex I/CS levels to lipid levels. We only found one significant lipid related to complex II-III levels – ceramide C22:6, and none significantly different in complex IV.

**Figure 6.**
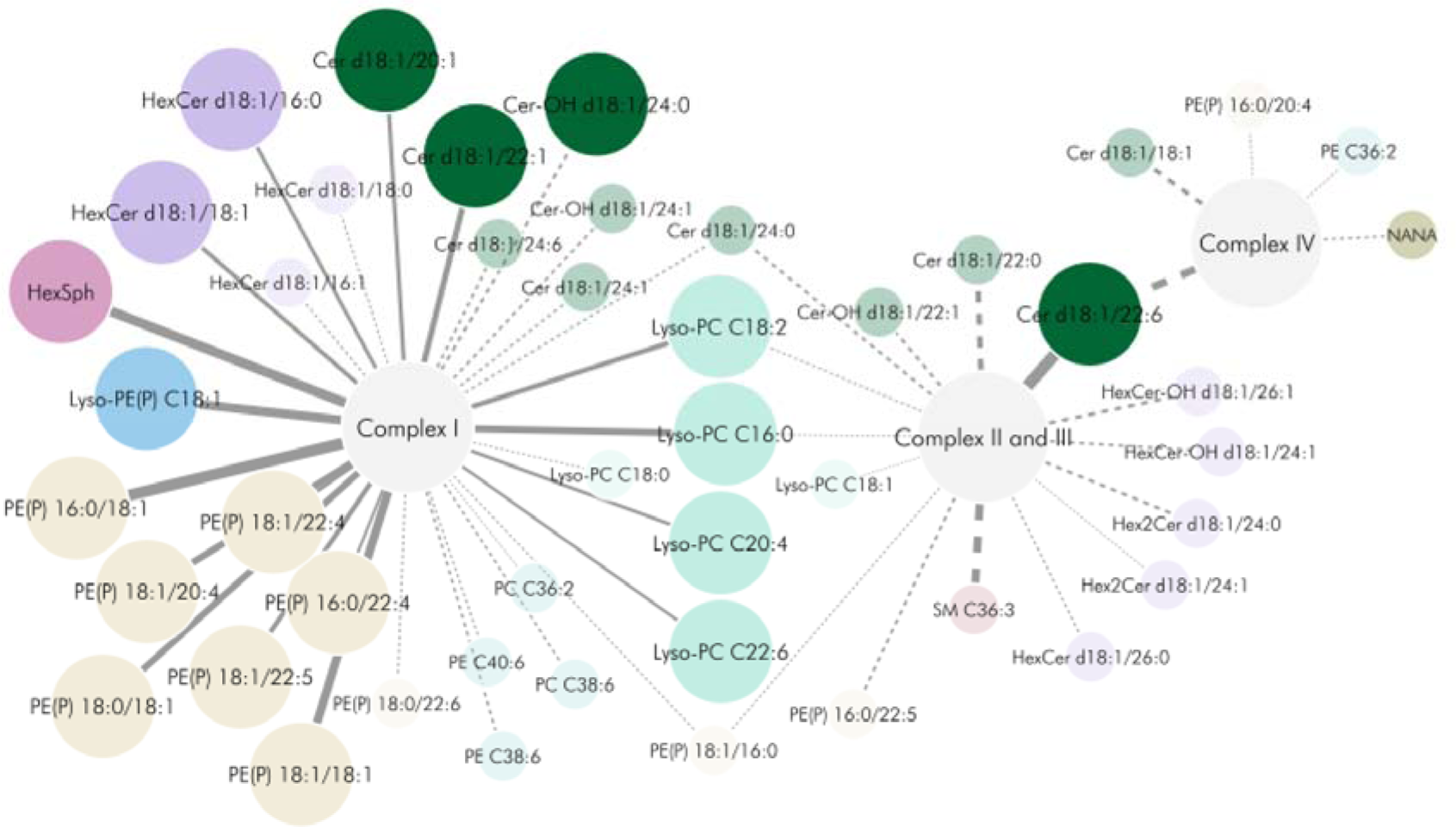
Network of linear mixed model results, showing significant interactions between class (PD or control), and mitochondrial complex I, II-III, IV ratioed to citrate synthase. The large circle nodes represent compounds statistically significant after Benjamini-Hochberg multiple testing correction at FDR = 10%, and the smaller radius nodes connected by dashed edges represent compounds significant on a nominal level.

The significant relationship between lipids and mitochondrial complex I suggests that these entities may modulate each other and would benefit from further investigation.

## Discussion

The spatial lipid profiling of anatomical brain regions revealed that each region possesses a unique lipid profile (**Figure 1A**). The regions subjected to lipid profiling included the deep brain nuclei area (putamen and caudate), the limbic area (parahippocampus and cingulate), the outer lobe regions (temporal, frontal, and parietal cortex), and the cerebellum. Each region exhibited a distinct lipid profile, although, apart from the outer lobes, the regions within the same areas did not exhibit similar lipid profiles. For instance, the caudate demonstrated more similarity with the parahippocampus than it did with its neighbouring deep brain nuclei region, the putamen (**Figure 2C** and **Figure 3D**).

The most conserved lipids, which exhibited minimal variation across brain regions, were ceramides. These had a coefficient of variation of 15% between brain regions, with only a higher normalised expression observed in the temporal cortex. This minimal variation in ceramide levels across the brain suggests that its regulation is critical, and its homeostasis is well controlled within the brain.

It is well-established that brain volume decreases with age and previous studies have highlighted the shrinkage of the hippocampus and cerebellum in particular^22^ with the dentate gyrus of the hippocampus being the region most susceptible to ageing^23^. Our analysis (Supplementary Results Figure S2) demonstrated that age has an evident effect on the brain lipid profile, with the putamen being strongest affected. Certain long-chain ceramide species, along with PC and lyso-PC, were strongly influenced by age, but it was n-hexosylceramide that exhibited the greatest overall change. These findings corroborate observations from previous studies in human and mice that have shown glucocerebrosidase activity progressively declining in the substantia nigra and the putamen in normal ageing, thus highlighting ageing as a risk factor for PD^24,25^. We did explore if changes in PD mirrored those of a non-pathological advanced aged brain, but the only similarities we found were elevation of sphingomyelin C34:2 and C34:3, phosphocholine C38:7 and lyso-SM 509 (an analogue of lyso-SM named *N*-palmitoyl-*O*-phosphocholine serine^26^ [PPCS]).

When analysing the overall changes occurring in the lipid profiles of sporadic PD to control by hierarchical clustering, we observed two clusters, one of which may represent the pattern of changes in early-stage PD, and one later which may correspond to late-stage PD disease spread. The two clusters demonstrated different changes, where the early affected regions showed increased levels of phospholipids and decreased levels of glycosphingolipids. The second cluster, representing regions affected in late-stage PD, showed an increase in gangliosides and some isoforms of HexCer and Hex2Cer, thereby indicating that the GSL pathway may be more relevant to these regions or that these are earlier events as these regions are latter affected.

It is noteworthy that the ceramides exhibited differential isoform changes, with an increase in long-chain ceramides (C14 and longer) and a decrease in very long-chain ceramides (C22 and longer). This differential effect has been previously observed in the brains of PD patients, along with an increase in ceramide synthase 1 (CERS1), which is responsible for generating the C18 isoforms^27^. The authors postulated that upregulation of CERS1 could be a downstream effect in response to stress in pathogenesis of PD. C18 ceramide has been found to induce mitophagy by anchoring LC3B II autophagolysosomes to mitochondrial membranes^28^. Mitophagy induced by accumulated ceramide has been described as an alternative pathway initiated by the cell to overcome defective PINK1 related mitophagy in PINK1 mutation PD. This mechanism of mitophagy may result in reduced β-oxidation leading to defective mitochondria, thereby increasing the need for clearance and resulting in a vicious cycle^29^. Our data and that of Abbot et al^27^ indicate that this alternative mechanism of mitophagy could be occurring in sporadic PD.

Gangliosides are complex glycosphingolipids and are situated at the beginning of the GSL degradation pathway (Supplementary Results Figure S1). Several studies have identified a reduction of gangliosides in the substantia nigra of PD patients^30,31^. This downregulation is believed to be specific to the region since the results from our analysis agrees with a recent study by Blumenreich et al^32^ showing a moderate elevation of gangliosides in sporadic PD. In likeness to Blumenreich et al, our study did not include the substantia nigra, the region most affected by PD, due to insufficient tissue amounts as a result of disease degeneration. The reason for the elevation of gangliosides in PD is unknown but could be caused by a stress-response effect as the ganglioside GM1, in particular, has neuroprotective and neurorestorative properties in Parkinsonism^33,34^ and has been trialled as a therapeutic in PD^35,36^.

When we divided the PD samples into early and late Braak stages in the three available regions and analysed these stages separately, we observed trends for lipids at the earliest stages of disease where the tissue remains pathologically unaffected by Lewy body disease (frontal cortex in Braak stage 3-4), or when the tissue has mild-moderate pathology (putamen in Braak stage 3-4). This suggests that alterations in lipid classes may underlie early pathogenic pathways in disease.

The cerebellum is often overlooked in PD studies in favour of the basal ganglia. However, there is increasing evidence that it does have significance in the pathophysiology of PD^37,38^. A recent study found that early-stage drug-naïve PD patients showed changes in cerebellar functional connectivity^39^ supporting its involvement at an early stage prior to clinical presentation of non-motor features of PD. Our findings indicate both early and transient involvement of the cerebellum, not just in the lipid profile but also in the proteome as previously described ^21^.

Plasmalogens are important in membrane dynamics and act as anti-oxidants^40^. They have previously been reported reduced in lipid rafts of early PD patients, with total levels in the brain unaffected^19^. We observed elevated levels of plasmalogens in the frontal cortex and putamen, consisting of a modest elevation in the frontal cortex of early- and late-stage PD, and a dramatic elevation in late-stage PD putamen. As stipulated for the gangliosides, this increase of plasmalogens could be providing a protective antioxidant effect.

Multi-omic analysis revealed that sphingomyelin correlated with mitochondrial proteins, but not with mitochondrial complex activity. However, we found that C18 ceramides were associated with complex I activity, along with C18 HexCer, C16-18 plasmalogens and lyso-PC. The C18 ceramide association corroborates that C18 induced mitophagy is occurring in early-stage PD. Other lipids also correlated with PD complex I activity, including plasmalogens and lyso-plasmalogen, which act as scavengers for reactive oxygen species. With this observation, their increase in PD brain tissue may be due to reactive oxygen species generated from complex I. Lyso-PC also correlated with complex I in PD brain tissue. This deacylated lipid is generated by phospholipase A2, an enzyme of which increased activity has been observed in the serum of PD patients^41^. Additionally, increased lyso-PC C16 and C18 have been identified previously in a rat model of early-stage PD, where the authors suggest that the increase could be caused by inflammatory processes. In relation to mitochondria, lyso-PC has been found to mildly affect mitochondrial permeability but significantly affect mitochondrial function^42^. Causation of sporadic PD has been found to be heterogenous for mitochondrial or lysosomal dysfunction^43^. This heterogeneity could confound the analysis; therefore, we examined the data for any subgrouping of lysosomal lipids to see if there was potential lysosomal dysfunction signature but could not identify any evident subgroups among the PD samples, thus suggesting that the dataset could be treated as a whole.

In conclusion, we describe a detailed lipid profile across the control human brain, and how these changes with anatomical region, age and disease. This will provide a resource for the community seeking to understand global lipid profiles in age related brain disorders. Our results highlight three key changes associated with Parkinson’s, namely (i) an elevation of long chain ceramides indicating mitophagy (ii) elevation of antioxidant lipids gangliosides and plasmalogens (iii) association of increased C34 and C36 sphingomyelin with proteomic changes.

## Materials and methods

### Brain tissue samples

The samples included in this study have been described by Toomey et al^21^. In short, post-mortem brain tissue was obtained from the Neurological Tissue Bank, IDIBAPS-HC-Biobanc, Barcelona; Human Brain Tissue Bank, Budapest; UK Parkinson’s Disease Society Tissue Bank, Imperial College London; the London Neurodegenerative Diseases Brain Bank, Institute of Psychiatry, King’s College, London; Netherlands Brain Bank, Amsterdam; and the Newcastle Brain Tissue Resource. Informed consent was given in all cases. Cases and controls were matched as close as possible for age and sex, and all had a post-mortem delay of less than 20 h. Control subjects did not have a diagnosis of neurological disease in life and at post-mortem displayed age-related changes only. All cases were assessed for alpha-synuclein pathology and rated according to Braak staging. PD cases were categorised as either Braak stage 3 or 4 (early) or Braak stage 6 (late). Ethical approval for the study was obtained from the Local Research Ethics Committee of the National Hospital for Neurology and Neurosurgery.

In total, we analysed 185 post-mortem biopsies from the brain regions caudate, cerebellum, cingulate cortex, frontal cortex, parahippocampus, parietal cortex, putamen and temporal cortex. At least 50% of both the early PD and the control samples were sampled in at least six of the regions. Late PD was generally only sampled in the frontal cortex, cerebellum and putamen. The demographics of the cohort are described in Supplementary Methods Table S1.

### Preparation of samples for targeted LC-MS/MS lipid analysis

The samples were prepared as described by Toomey et al^21^. In brief, the post-mortem brain tissue was homogenised and an amount of homogenate equivalent to 300 mg of total protein was precipitated with acetone. The lipid-containing supernatant was collected, and evaporated using a vacuum concentrator, after which the dry metabolite fraction was stored at -80 °C until further use. The lipid-containing fraction was reconstituted in 75 µL methanol containing the internal standards d_3_-ceramide (Cayman chemicals, 24396), d_3_-GM1 (Matreya, 2050), d_3_-GM2 (Matreya, 2051), d_3_-GM3 (Matreya, 2052), d_5_-glucosylsphingosine (Avanti polar lipids, 860636), d_3_-glucosylceramide (Matreya, 1533) and N-glycinated globotriaosylsphingosine (Matreya, 1530). The samples were shaken on a rotational shaker at 1500 rpm for 15 minutes and sonicated for 15 minutes prior to centrifugation at 16 900 x *g* for five minutes. 70 µL supernatant was transferred to glass micro vials.

### Analysis of lipids by targeted LC-MS/MS

In total, we quantified 135 species belonging to 11 main compound classes. The samples were analysed using a triple quadrupole mass spectrometer (Waters TQ-S) equipped with an ESI source, coupled to a quaternary Waters Acquity liquid chromatographic separation system. Detection was performed in multiple reaction monitoring mode (MRM), where the transitions from precursor ions to class-specific fragment ions were monitored. The LC methods were tailored to achieve maximum separation between the compound classes. To ensure that adequate analytical setups were employed for each of the compound classes, we utilised three separate LC-MS/MS methods with different column chemistries (HILIC, Amide, or C8). Details about the chromatographic separation parameters can be found in **Supplementary Methods Table S2** and details about the multiple reaction monitoring (MRM) transitions, MS settings and ionisation modes can be found in **Supplementary Methods Table S3**. The compound classes were normalised to internal standards according to **Table 1**.

### Data integration

Data were acquired in MassLynx 4.2 and transformed into text files using the application MSConvert from the package ProteoWizard^44^. Peak picking and integrations were performed using an in-house application written in Python (available via the GitHub repository https://github.com/jchallqvist/mrmIntegrate) which rendered area under the curve by the trapezoidal integration method. Each analyte was thereafter normalised to an internal standard as described in **Table 1**.

### Statistical methods and visualisation

If not otherwise specified, all statistical analyses were performed in Python (version 3.8.16). The dataset was inspected for outliers and instrumental drift using principal component analysis (PCA) and orthogonal projection to latent variables (OPLS) in SIMCA, version 17 (Umetrics Sartorius Stedim, Umeå, Sweden). Outliers exceeding ten median absolute deviations from each variable’s median were excluded. Adjustment for age and regions was performed using multiple linear regression from Statsmodels (version 0.13.5). The data were evaluated for normal distribution using D’Angostino and Pearson’s method from SciPy (version 1.9.3). Significance testing between the independent groups of control and PD, and of control, early-stage PD and late-stage PD, was performed by Student’s two-tailed t-test for normally distributed variables and by Mann-Whitney’s non-parametric U-test, both from the SciPy’s stats package, for the non-normally distributed variables. The Benjamini-Hochberg FDR discovery rate procedure (Statsmodels version 0.14.0) was applied with alpha set to 0.1. Fold-changes were calculated by dividing the means or medians of the affected groups by the control group. Correlation analyses in the targeted data were performed by Spearman’s correlation (SciPy) and the correlation p-values were adjusted variable-wise by the Benjamini-Hochberg FDR procedure. UMAP projections were performed using the Python package Umap (version 0.5.3)^20^. Box plots and hierarchical clusters were produced using the Seaborn (version 0.12.2) and Matplotlib (version 3.7.0) libraries. Linear mixed models were performed using the R-to-Python bridge software pymer4^45^ (version 0.8.0), where region was set as a random effect and the interaction between the classes PD/control and the mitochondrial complexes I, II-III and IV were evaluated for significance post Benjamini-Hochberg’s procedure^46^ for multiple testing correction.

All multivariate analyses were performed in SIMCA. OPLS and OPLS-discriminant analysis (OPLS-DA) models were evaluated for significance by ANOVA p-values and by permutation tests applying 1000 permutations, where p < 0.05 and p < 0.001 were deemed significant, respectively.

Data were analysed for pathway enrichment and Gene Ontology (GO) annotations using DAVID Bioinformatics Resources (2021 build). Networks were built in Cytoscape (version 3.8.0) applying the “Organic layout” from yFiles.

## Ethics

Ethical approval for the study was obtained from the Local Research Ethics Committee of the National Hospital for Neurology and Neurosurgery, Queen Square, London.

## Acknowledgements

The authors gratefully acknowledge support and funding from the Michael J Fox Foundation, Parkinson’s UK, the Peto Foundation, the Translational Mass Spectrometry Research Group at UCL, Eisai the Lila Reta Weston Trust and Aligning Science Across Parkinson’s. This work is supported by the NIHR GOSH BRC. The views expressed are those of the authors and not necessarily those of the NHS, the NIHR or the Department of Health. RP is supported by the UK Dementia Research Institute (award number MC_PC_17114), which receives its funding from UK Dementia Research Institute Ltd., funded by the UK Medical Research Council (MRC), Alzheimer’s Society, and Alzheimer’s Research UK.

## Authors’ contributions

JH: Conceptualization, Data acquisition, Data curation, Formal analysis, Methodology, Writing – original draft. CET: Data acquisition, Data curation, Formal analysis, Writing – review & editing. RP: Formal Analysis, Methodology, Writing – review & editing. AW, MAS: Data acquisition, Formal analysis SE, SH: Supervision – review & editing. KM, SG and WEH: Conceptualization, Supervision, Methodology, Writing – original draft.

## Competing interests

None of the authors have any competing interests to report.

## Data availability

The raw targeted chromatograms of the lipid data are available to view and download via the Panorama repository (https://panoramaweb.org/PD_Brain_Lipids.url).

## Notes

### Competing Interest Statement

The authors have declared no competing interest.

