## Supplementary Figures, tables and methods for "Multi-Omic Analysis Reveals Lipid Dysregulation Associated with Mitochondrial Dysfunction in Parkinson’s Disease Brain"

**Supplementary Results**

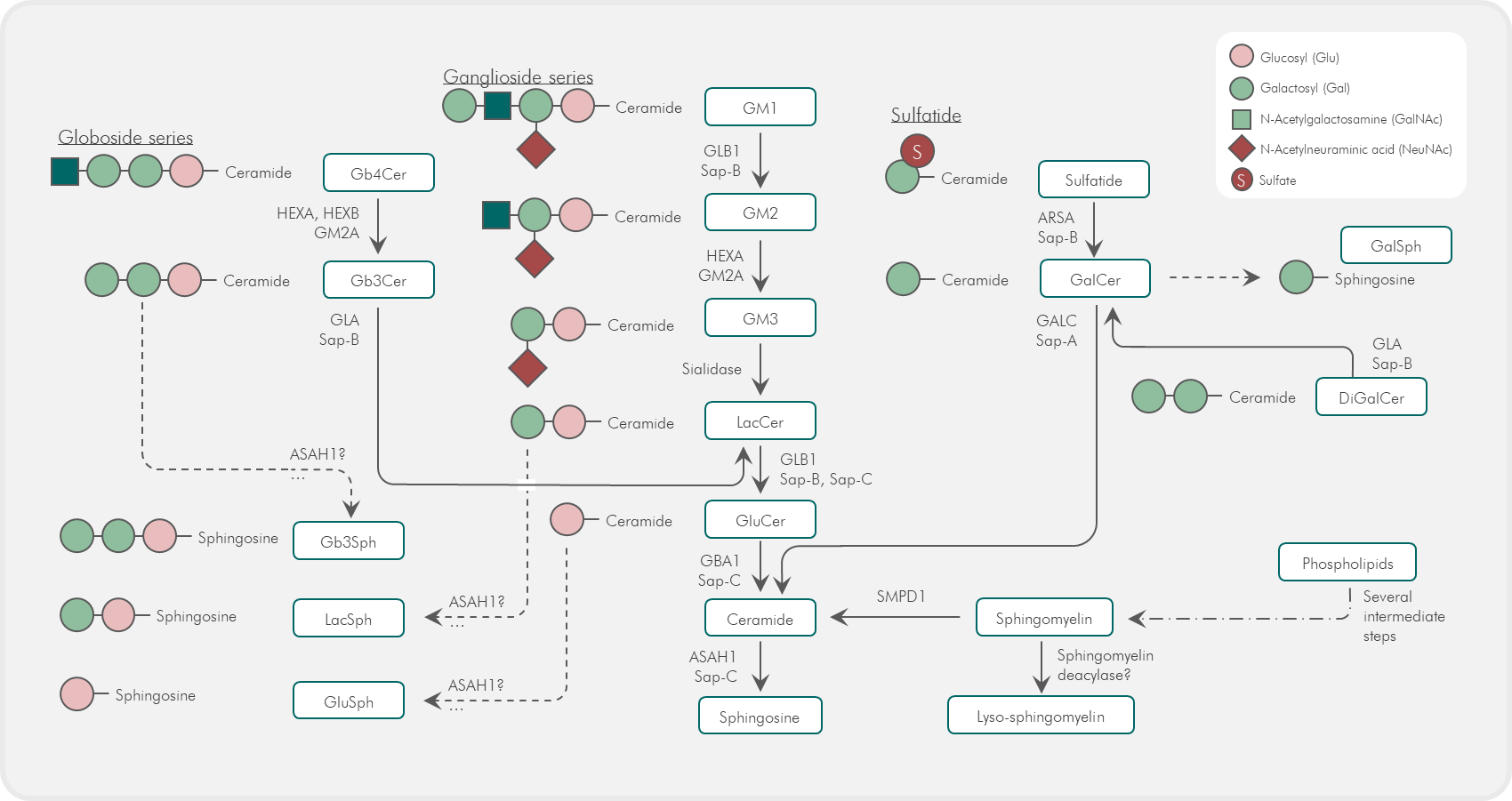

**Supplementary Results Figure S1.** **The catabolic lysosomal glycosphingolipid degradation pathway.** *The pathway shows lipids from the globoside, ganglioside and sulfatide series and the enzymes catabolising each step. The diseases associated with malfunctioning of the enzymes are also shown.*

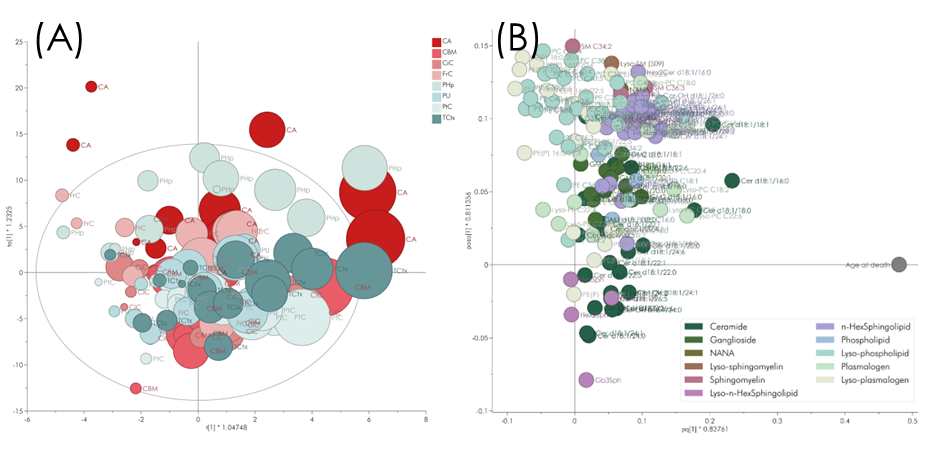

**Supplementary Results Figure S2.** **Relationship between age and lipids in control samples.** *(A) OPLS scores from the evaluation of the correlation between age and lipids in controls, sized by age. The OPLS model with age set as the dependent variable was found significant with ANOVA p = 1.9 E^-6^ and permutations p << 0.001 thereby indicating an age effect on the brain lipid levels. (B) OPLS loadings from the evaluation of the correlation between age and lipids in controls. Ceramides with C16 and C18 fatty acid chains showed the strongest correlation with increasing age, along with several n-hexosylceramide species.*

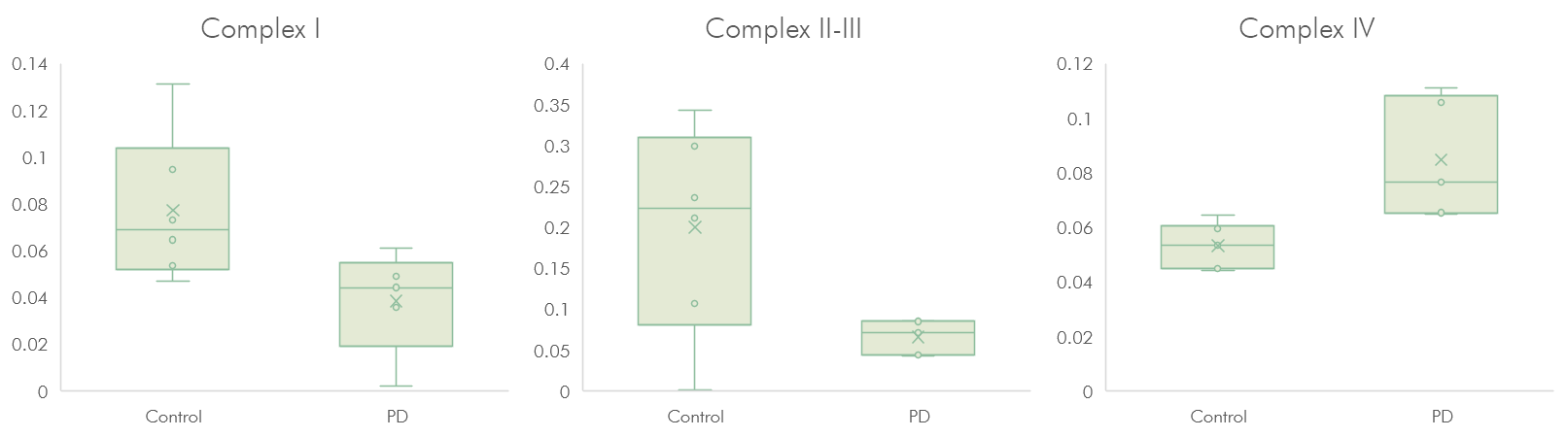

**Supplementary Results Figure S3.** **Ratios of complexes I, II-III and IV to citrate synthase in the putamen region.** *All the complex ratios demonstrated significant differences between PD and control.*

**Supplementary Results Table S1.** **p-values showing if the slope of a linear regression of lipid levels versus age was significantly non-zero.** *From the region-wise linear regression of lipids and age at the time of death in all samples. The significantly non-zero slopes are highlighted in green and were age-adjusted. Remaining and non-affected compounds were not age-adjusted.*

|  | **Caudate** | **Cerebellum** | **Cingulate cortex** | **Frontal cortex** | **Parahippo-campus** | **Parietal cortex** | **Putamen** | **Temporal cortex** |
| --- | --- | --- | --- | --- | --- | --- | --- | --- |
| Cer d18:1/16:0 | 1.5E-01 | 3.4E-02 | 5.4E-02 | 4.7E-01 | 4.8E-03 | 2.6E-04 | 3.6E-01 | 1.4E-01 |
| Cer d18:1/16:1 | 4.3E-01 | 9.3E-04 | 6.7E-02 | 5.2E-01 | 1.3E-01 | 2.6E-01 | 3.2E-01 | 1.8E-01 |
| Cer d18:1/18:0 | 4.0E-02 | 3.5E-02 | 1.8E-01 | 4.3E-01 | 7.9E-02 | 2.0E-01 | 9.4E-01 | 5.6E-01 |
| Cer d18:1/18:1 | 1.8E-01 | 1.9E-03 | 1.3E-03 | 1.6E-01 | 1.9E-02 | 2.2E-02 | 3.2E-01 | 1.5E-03 |
| Cer d18:1/20:0 | 4.6E-01 | 8.7E-01 | 7.5E-01 | 9.0E-01 | 5.7E-01 | 4.2E-01 | 4.9E-01 | 7.2E-01 |
| Cer d18:1/20:1 | 4.3E-01 | 2.3E-02 | 5.9E-01 | 1.0E+00 | 3.0E-01 | 2.0E-02 | 8.0E-01 | 8.9E-01 |
| Cer d18:1/20:4 | 4.0E-01 | 2.6E-01 | 8.1E-01 | 3.8E-01 | 1.7E-01 | 8.0E-01 | 9.5E-01 | 6.2E-01 |
| Cer d18:1/20:5 | 1.8E-01 | 7.4E-02 | 8.2E-01 | 7.8E-01 | 8.7E-01 | 7.3E-01 | 6.9E-01 | 3.6E-01 |
| Cer d18:1/22:0 | 5.8E-01 | 2.1E-01 | 5.7E-01 | 4.2E-01 | 5.3E-01 | 1.7E-01 | 4.0E-01 | 4.1E-01 |
| Cer d18:1/22:1 | 6.6E-01 | 6.0E-02 | 6.7E-01 | 5.6E-01 | 5.8E-01 | 2.6E-02 | 9.9E-01 | 9.0E-01 |
| Cer d18:1/22:5 | 5.2E-01 | 4.7E-01 | 8.8E-01 | 6.3E-01 | 9.7E-01 | 2.7E-01 | 1.2E-01 | 9.2E-01 |
| Cer d18:1/22:6 | 9.9E-01 | 1.3E-02 | 6.6E-02 | 2.7E-01 | 4.9E-01 | 6.9E-02 | 9.3E-01 | 9.7E-01 |
| Cer d18:1/24:0 | 8.0E-01 | 3.1E-02 | 6.0E-01 | 5.7E-01 | 9.3E-01 | 3.1E-02 | 2.8E-01 | 4.9E-01 |
| Cer d18:1/24:1 | 8.1E-01 | 9.9E-02 | 4.1E-01 | 5.9E-01 | 7.4E-01 | 3.3E-02 | 4.3E-01 | 7.2E-01 |
| Cer d18:1/24:2 | 8.0E-01 | 2.7E-01 | 6.7E-01 | 7.5E-01 | 7.5E-01 | 4.7E-02 | 9.9E-01 | 7.2E-01 |
| Cer d18:1/24:6 | 5.0E-01 | 1.3E-01 | 5.5E-01 | 1.9E-01 | 9.1E-01 | 3.2E-01 | 2.8E-01 | 2.0E-01 |
| Cer d18:1/26:5 | 5.1E-01 | 5.3E-01 | 3.4E-01 | 8.1E-01 | 2.2E-01 | 7.2E-01 | 8.1E-01 | 2.4E-01 |
| Cer d18:1/26:6 | 3.2E-01 | 8.8E-03 | 4.3E-01 | 8.5E-01 | 8.4E-01 | 2.4E-02 | 4.9E-01 | 2.5E-01 |
| Cer-OH d18:1/16:0 | 7.3E-01 | 8.9E-02 | 4.1E-01 | 6.3E-01 | 8.6E-01 | 4.2E-01 | 5.4E-01 | 7.3E-01 |
| Cer-OH d18:1/18:0 | 3.0E-01 | 1.2E-01 | 4.1E-01 | 6.3E-01 | 2.1E-02 | 4.1E-01 | 8.3E-01 | 8.5E-01 |
| Cer-OH d18:1/22:1 | 7.0E-01 | 2.3E-01 | 4.9E-01 | 7.4E-01 | 8.9E-01 | 4.2E-02 | 1.8E-01 | 5.7E-01 |
| Cer-OH d18:1/24:0 | 4.9E-01 | 3.2E-03 | 4.7E-01 | 6.2E-01 | 5.4E-01 | 6.1E-02 | 1.5E-01 | 5.7E-01 |
| Cer-OH d18:1/24:1 | 4.2E-01 | 2.2E-02 | 3.1E-01 | 7.5E-01 | 2.0E-01 | 1.1E-01 | 5.7E-01 | 5.5E-01 |
| GM3 d18:1/18:0 | 3.3E-01 | 9.7E-01 | 3.3E-02 | 2.7E-01 | 6.2E-01 | 5.5E-02 | 1.6E-01 | 2.7E-02 |
| GM2 d18:1/16:0 | 7.4E-01 | 3.6E-01 | 1.9E-01 | 8.5E-02 | 6.7E-02 | 3.6E-01 | 1.5E-01 | 4.2E-04 |
| GM2 d18:1/18:0 | 6.0E-01 | 3.7E-01 | 1.5E-01 | 7.3E-01 | 9.1E-01 | 1.6E-01 | 4.9E-01 | 3.8E-02 |
| GM2 d18:1/18:1 | 2.8E-01 | 1.1E-01 | 7.1E-02 | 3.6E-01 | 3.9E-01 | 3.1E-01 | 4.7E-01 | 1.1E-02 |
| GM2 d18:1/20:0 | 6.0E-01 | 6.3E-01 | 8.5E-02 | 7.5E-01 | 8.6E-01 | 5.0E-01 | 3.2E-01 | 5.9E-02 |
| GM1 d18:1/16:0 | 8.8E-01 | 4.2E-01 | 1.3E-01 | 1.8E-01 | 5.9E-01 | 5.1E-01 | 1.2E-01 | 2.4E-03 |
| GM1 d18:1/18:0 | 7.3E-01 | 4.6E-01 | 1.3E-01 | 7.2E-01 | 3.3E-02 | 1.4E-01 | 3.9E-01 | 1.3E-02 |
| GM1 d18:1/18:1 | 4.0E-01 | 1.4E-01 | 1.2E-01 | 4.4E-01 | 4.2E-01 | 4.2E-01 | 1.1E-01 | 7.4E-03 |
| GM1 d18:1/20:0 | 6.2E-01 | 6.5E-01 | 7.9E-02 | 9.2E-01 | 5.6E-02 | 6.7E-01 | 4.5E-01 | 1.4E-02 |
| GM1 d18:1/20:1 | 3.2E-01 | 2.3E-01 | 1.2E-01 | 7.8E-01 | 4.1E-01 | 3.2E-01 | 2.3E-01 | 9.8E-03 |
| GM1 d18:1/22:0 | 5.2E-01 | 7.8E-01 | 1.5E-01 | 7.7E-01 | 6.5E-01 | 3.6E-01 | 4.0E-01 | 1.2E-01 |
| GM1 d18:1/24:1 | 2.5E-01 | 2.3E-01 | 2.7E-01 | 4.4E-01 | 2.2E-02 | 4.5E-01 | 4.4E-01 | 2.5E-02 |
| NANA | 4.3E-01 | 2.0E-01 | 3.9E-01 | 4.1E-01 | 3.9E-01 | 3.2E-01 | 4.8E-02 | 2.5E-01 |
| Lyso-SM C16:1 | 3.2E-01 | 3.5E-02 | 6.2E-04 | 3.1E-01 | 8.1E-02 | 2.1E-01 | 3.2E-01 | 2.2E-01 |
| Lyso-SM C18:1 | 8.9E-01 | 4.1E-01 | 1.0E-01 | 1.1E-01 | 9.9E-02 | 5.7E-01 | 4.3E-01 | 1.9E-01 |
| Lyso-SM (509) | 5.0E-01 | 1.9E-01 | 6.4E-01 | 2.3E-01 | 9.4E-02 | 4.0E-01 | 5.6E-02 | 1.1E-01 |
| SM C34:2 | 1.2E-01 | 5.1E-01 | 6.0E-01 | 7.7E-01 | 6.8E-01 | 9.1E-01 | 1.6E-01 | 7.0E-02 |
| SM C34:3 | 2.6E-01 | 2.0E-01 | 9.9E-03 | 3.2E-01 | 4.9E-02 | 6.8E-02 | 1.1E-01 | 1.3E-03 |
| SM C36:3 | 5.5E-01 | 8.1E-02 | 1.2E-02 | 2.7E-01 | 4.4E-02 | 1.6E-01 | 1.7E-01 | 3.1E-04 |
| HexSph -28 | 4.0E-01 | 7.1E-01 | 1.1E-01 | 3.7E-01 | 5.6E-01 | 8.3E-01 | 4.8E-02 | 3.3E-02 |
| HexSph | 8.8E-01 | 8.2E-02 | 9.6E-01 | 9.3E-01 | 9.5E-01 | 6.9E-02 | 4.4E-01 | 4.7E-01 |
| HexCer d18:1/16:1 | 4.1E-01 | 1.5E-01 | 6.1E-01 | 3.0E-01 | 1.7E-01 | 5.7E-01 | 7.3E-01 | 1.8E-01 |
| HexCer d18:1/16:0 | 7.5E-01 | 2.2E-01 | 5.4E-01 | 9.1E-01 | 2.2E-01 | 3.0E-02 | 7.8E-01 | 8.0E-01 |
| HexCer d18:1/18:1 | 6.0E-01 | 1.6E-01 | 3.6E-01 | 8.6E-01 | 5.0E-01 | 1.9E-02 | 4.9E-01 | 6.5E-01 |
| HexCer d18:1/18:0 | 9.5E-03 | 6.0E-01 | 8.0E-01 | 2.5E-01 | 4.8E-01 | 2.8E-01 | 2.0E-01 | 9.5E-01 |
| HexCer-OH d18:1/18:0 | 3.4E-01 | 6.0E-01 | 4.6E-01 | 2.5E-01 | 6.3E-02 | 1.5E-01 | 1.8E-01 | 9.9E-03 |
| HexCer-OH d18:1/20:0 | 7.1E-01 | 6.7E-01 | 7.8E-01 | 2.5E-01 | 5.3E-02 | 2.5E-01 | 3.7E-01 | 9.5E-03 |
| HexCer-OH d18:1/22:0 | 3.2E-01 | 6.0E-01 | 2.8E-01 | 3.7E-01 | 2.4E-01 | 1.7E-02 | 4.0E-01 | 1.1E-01 |
| HexCer d18:1/24:1 | 2.5E-01 | 4.1E-01 | 4.5E-01 | 2.3E-01 | 1.9E-01 | 4.9E-02 | 1.5E-01 | 1.4E-01 |
| HexCer d18:1/24:0 | 6.6E-01 | 8.6E-01 | 6.4E-01 | 2.8E-01 | 1.9E-01 | 1.4E-01 | 5.3E-01 | 4.8E-02 |
| HexCer-OH d18:1/24:2 | 2.3E-01 | 4.0E-01 | 3.4E-01 | 2.6E-01 | 8.6E-02 | 6.0E-03 | 1.2E-01 | 1.6E-02 |
| HexCer-OH d18:1/24:1 | 4.4E-01 | 3.1E-01 | 2.8E-01 | 1.4E-01 | 1.2E-01 | 2.8E-04 | 4.8E-02 | 2.4E-03 |
| HexCer-OH d18:1/24:0 | 7.9E-01 | 8.5E-01 | 1.9E-01 | 3.5E-01 | 1.8E-01 | 7.6E-04 | 1.2E-01 | 4.0E-03 |
| HexCer d18:1/26:1 | 2.6E-01 | 6.0E-01 | 5.0E-01 | 2.8E-01 | 5.7E-02 | 1.6E-01 | 1.9E-01 | 1.4E-02 |
| HexCer d18:1/26:0 | 4.2E-01 | 2.1E-01 | 1.7E-01 | 2.3E-01 | 5.7E-02 | 3.5E-04 | 3.0E-02 | 2.2E-03 |
| HexCer-OH d18:1/26:1 | 3.0E-01 | 2.1E-01 | 1.7E-01 | 1.5E-01 | 1.0E-01 | 6.1E-04 | 3.4E-02 | 4.3E-03 |
| HexCer-OH d18:1/26:0 | 7.0E-01 | 5.5E-01 | 1.4E-01 | 3.5E-01 | 1.2E-01 | 5.3E-03 | 3.5E-01 | 1.2E-03 |
| Hex2Sph | 6.5E-01 | 2.0E-01 | 5.1E-01 | 7.7E-01 | 2.7E-01 | 9.8E-01 | 2.6E-01 | 3.1E-01 |
| Hex2Cer d18:1/16:0 | 2.1E-01 | 5.8E-01 | 5.3E-01 | 5.5E-01 | 9.4E-03 | 8.8E-01 | 4.1E-01 | 1.3E-01 |
| Hex2Cer d18:1/18:0 | 1.7E-01 | 8.3E-01 | 1.5E-01 | 3.3E-01 | 1.3E-03 | 6.9E-01 | 2.9E-01 | 4.4E-02 |
| Hex2Cer d18:1/24:1 | 2.0E-01 | 7.7E-01 | 1.8E-01 | 2.1E-01 | 3.9E-03 | 1.7E-02 | 5.6E-03 | 4.8E-03 |
| Hex2Cer d18:1/24:0 | 5.6E-01 | 8.5E-01 | 1.2E-01 | 3.5E-01 | 2.7E-03 | 1.0E-02 | 3.4E-02 | 3.0E-03 |
| Hex2Cer-OH d18:1/24:2 | 2.4E-01 | 5.4E-01 | 5.8E-02 | 1.6E-01 | 4.5E-03 | 1.1E-02 | 1.3E-02 | 4.1E-03 |
| Hex2Cer-OH d18:1/24:1 | 9.3E-01 | 4.1E-01 | 1.1E-01 | 2.1E-01 | 2.6E-03 | 3.2E-02 | 8.3E-02 | 7.2E-04 |
| Gb4Cer d18:1/16:0 | 3.5E-01 | 5.5E-01 | 2.6E-02 | 9.5E-01 | 3.8E-01 | 8.1E-02 | 6.9E-02 | 4.0E-02 |
| Gb3Sph | 8.5E-01 | 4.6E-01 | 5.0E-01 | 7.0E-01 | 7.5E-01 | 7.1E-02 | 4.7E-02 | 1.3E-01 |
| Gb3Cer d18:1/16:0 | 2.5E-01 | 8.1E-01 | 1.1E-01 | 4.9E-01 | 2.5E-01 | 5.9E-02 | 1.8E-01 | 6.1E-02 |
| Gb3Cer d18:1/18:0 | 8.3E-01 | 5.4E-02 | 3.2E-01 | 7.8E-01 | 9.5E-02 | 6.0E-01 | 3.3E-01 | 7.5E-01 |
| Lyso-PE(P) C16:0 | 2.2E-01 | 3.9E-01 | 9.9E-01 | 3.8E-01 | 2.7E-01 | 4.2E-01 | 4.5E-01 | 2.0E-01 |
| Lyso-PE C16:0 | 1.6E-01 | 1.1E-01 | 4.0E-01 | 6.2E-01 | 8.2E-01 | 8.7E-01 | 8.3E-01 | 5.0E-01 |
| Lyso-PE(P) C18:1 | 9.8E-01 | 7.2E-02 | 8.3E-01 | 2.3E-01 | 5.1E-01 | 1.3E-01 | 1.9E-01 | 3.0E-01 |
| Lyso-PE(P) C18:0 | 1.1E-01 | 5.2E-01 | 3.1E-01 | 4.4E-01 | 7.5E-02 | 2.4E-01 | 2.3E-01 | 4.8E-02 |
| Lyso-PE C18:2 | 7.8E-01 | 1.2E-01 | 4.6E-01 | 3.5E-01 | 5.1E-01 | 5.3E-01 | 4.6E-01 | 9.6E-02 |
| Lyso-PE C18:1 | 7.6E-01 | 6.6E-02 | 7.0E-01 | 4.3E-01 | 8.3E-01 | 5.7E-01 | 8.2E-01 | 4.9E-01 |
| Lyso-PC C16:0 | 5.8E-01 | 2.9E-01 | 5.0E-02 | 3.9E-01 | 4.8E-02 | 7.9E-01 | 5.0E-01 | 1.2E-01 |
| Lyso-PE C20:4 | 4.1E-01 | 3.1E-01 | 9.5E-01 | 9.4E-02 | 7.8E-01 | 4.2E-01 | 9.1E-01 | 8.7E-01 |
| Lyso-PC C18:2 | 5.0E-01 | 7.1E-01 | 1.4E-01 | 8.5E-01 | 3.0E-01 | 3.4E-01 | 5.2E-01 | 2.6E-01 |
| Lyso-PC C18:1 | 2.4E-01 | 1.1E-01 | 4.0E-02 | 5.3E-01 | 1.9E-01 | 6.6E-01 | 3.9E-01 | 4.8E-01 |
| Lyso-PC C18:0 | 7.4E-01 | 5.2E-01 | 1.0E-01 | 3.9E-01 | 2.2E-01 | 4.1E-01 | 2.0E-01 | 5.3E-02 |
| Lyso-PE C22:6 | 7.1E-01 | 4.0E-01 | 8.6E-02 | 5.2E-01 | 5.8E-01 | 8.4E-02 | 6.4E-01 | 3.9E-01 |
| Lyso-PE C22:5 | 4.8E-01 | 3.7E-01 | 5.9E-01 | 2.3E-01 | 5.6E-01 | 7.4E-01 | 7.5E-01 | 1.7E-01 |
| Lyso-PE C22:4 | 1.6E-02 | 3.0E-01 | 1.7E-01 | 9.1E-02 | 4.4E-02 | 3.3E-01 | 5.4E-01 | 3.9E-01 |
| Lyso-PC C20:4 | 3.3E-01 | 4.5E-01 | 6.6E-02 | 9.6E-01 | 2.5E-01 | 9.3E-01 | 6.8E-01 | 2.6E-01 |
| Lyso-PC C22:6 | 1.6E-01 | 2.2E-01 | 5.8E-03 | 3.2E-01 | 2.8E-01 | 5.0E-02 | 6.3E-02 | 2.1E-01 |
| Lyso-PC C22:5 | 1.5E-01 | 1.8E-01 | 4.9E-02 | 4.9E-01 | 2.1E-01 | 4.8E-01 | 5.3E-02 | 1.4E-01 |
| Lyso-PC C22:4 | 7.1E-01 | 2.6E-01 | 1.3E-01 | 7.9E-01 | 9.2E-01 | 4.5E-01 | 7.9E-01 | 5.7E-01 |
| PE C34:2 | 2.8E-01 | 1.3E-01 | 1.5E-01 | 7.8E-01 | 3.4E-02 | 6.4E-01 | 7.8E-01 | 6.3E-01 |
| PE C34:1 | 1.5E-01 | 4.4E-01 | 6.3E-01 | 4.0E-01 | 5.2E-02 | 6.6E-01 | 9.2E-01 | 8.0E-01 |
| PE C36:5 | 9.1E-01 | 6.2E-01 | 1.3E-01 | 1.0E+00 | 5.2E-02 | 9.7E-01 | 5.7E-01 | 5.7E-01 |
| PE C36:4 | 4.2E-01 | 3.9E-01 | 5.9E-01 | 3.1E-01 | 8.3E-01 | 9.5E-01 | 7.8E-01 | 7.2E-01 |
| PE C36:3 | 3.2E-01 | 7.5E-01 | 7.6E-02 | 3.1E-01 | 3.8E-01 | 4.4E-01 | 8.1E-01 | 7.0E-01 |
| PE C36:2 | 1.1E-01 | 1.9E-01 | 6.3E-01 | 6.6E-01 | 5.7E-01 | 1.2E-01 | 5.1E-01 | 4.4E-01 |
| PE C36:1 | 9.5E-01 | 6.4E-01 | 8.3E-01 | 2.7E-01 | 2.7E-01 | 3.4E-01 | 5.2E-01 | 4.2E-01 |
| PE(P) 18:0/20:4 | 5.7E-02 | 8.6E-01 | 7.4E-01 | 2.1E-01 | 6.7E-01 | 8.3E-01 | 7.7E-01 | 6.3E-01 |
| PE(P) 16:0/22:4 | 7.3E-02 | 6.3E-01 | 1.8E-01 | 2.4E-01 | 5.3E-01 | 6.1E-02 | 9.9E-01 | 6.8E-01 |
| PE C38:7 | 8.5E-01 | 5.4E-01 | 3.5E-02 | 6.5E-01 | 1.9E-01 | 6.1E-01 | 2.9E-01 | 2.7E-01 |
| PE C38:6 | 7.2E-01 | 6.0E-01 | 3.5E-02 | 7.7E-01 | 9.7E-01 | 6.1E-01 | 4.6E-01 | 6.0E-01 |
| PE C38:5 | 6.5E-01 | 7.3E-01 | 1.9E-01 | 6.0E-01 | 7.4E-01 | 6.7E-01 | 5.2E-01 | 6.4E-01 |
| PE C38:4 | 2.6E-01 | 8.1E-01 | 3.0E-01 | 4.1E-01 | 9.7E-01 | 9.6E-01 | 3.9E-01 | 5.8E-01 |
| PE(P) 18:1/22:6 | 8.9E-01 | 4.9E-02 | 2.1E-02 | 1.9E-01 | 2.8E-01 | 3.1E-01 | 1.8E-01 | 1.4E-01 |
| PE(P) 18:0/22:6 | 4.7E-01 | 7.6E-01 | 6.8E-02 | 5.9E-01 | 4.5E-01 | 3.3E-01 | 2.2E-01 | 1.9E-01 |
| PE(P) 18:1/22:5 | 8.0E-01 | 6.7E-02 | 8.3E-01 | 8.3E-01 | 3.9E-01 | 9.4E-02 | 5.9E-01 | 8.4E-01 |
| PE(P) 18:0/22:5 | 2.6E-01 | 8.9E-01 | 5.9E-01 | 6.7E-01 | 9.6E-01 | 5.4E-01 | 4.4E-01 | 3.5E-01 |
| PE(P) 18:1/22:4 | 8.6E-01 | 1.7E-01 | 3.5E-01 | 9.8E-01 | 9.2E-01 | 3.1E-02 | 9.2E-01 | 7.3E-01 |
| PE(P) 18:0/22:4 | 6.4E-02 | 8.7E-01 | 8.6E-01 | 3.0E-01 | 8.2E-01 | 7.1E-01 | 7.1E-01 | 3.8E-01 |
| PE C40:8 | 9.0E-01 | 9.4E-01 | 1.9E-01 | 8.7E-01 | 5.6E-01 | 7.7E-01 | 4.1E-01 | 8.8E-01 |
| PE C40:7 | 9.9E-01 | 6.6E-01 | 1.5E-02 | 4.1E-01 | 3.7E-01 | 6.6E-01 | 3.2E-01 | 3.3E-01 |
| PE C40:6 | 6.9E-01 | 8.8E-01 | 5.5E-03 | 5.9E-01 | 4.6E-01 | 6.2E-01 | 3.9E-01 | 4.5E-01 |
| PE C40:5 | 4.9E-01 | 7.9E-01 | 9.6E-02 | 7.4E-01 | 8.7E-01 | 8.0E-01 | 6.3E-01 | 7.2E-01 |
| PE C40:4 | 1.4E-01 | 5.6E-01 | 3.9E-01 | 3.5E-01 | 6.2E-01 | 9.4E-01 | 7.5E-01 | 8.1E-01 |
| PE(P) 18:1/16:0 | 5.9E-01 | 2.6E-01 | 3.1E-01 | 4.8E-01 | 4.8E-01 | 2.4E-01 | 5.2E-01 | 8.8E-01 |
| PE(P) 16:0/18:1 | 9.1E-01 | 9.7E-01 | 2.3E-01 | 8.5E-01 | 7.1E-01 | 1.0E-01 | 2.2E-01 | 6.5E-01 |
| PE(P) 16:0/20:4 | 7.0E-02 | 2.8E-01 | 2.3E-01 | 6.3E-02 | 9.6E-01 | 3.5E-01 | 6.5E-01 | 8.8E-01 |
| PE(P) 18:1/18:1 | 6.3E-01 | 2.1E-01 | 5.9E-01 | 6.8E-01 | 8.0E-01 | 4.2E-02 | 4.7E-01 | 7.4E-01 |
| PE(P) 18:0/18:1 | 4.7E-01 | 7.3E-01 | 1.7E-01 | 5.7E-01 | 6.2E-01 | 1.1E-01 | 3.1E-01 | 8.6E-01 |
| PE(P) 16:0/22:6 | 5.5E-01 | 7.3E-01 | 4.4E-01 | 4.5E-01 | 8.8E-01 | 4.5E-01 | 1.6E-01 | 1.6E-01 |
| PE(P) 18:1/20:4 | 5.6E-01 | 5.8E-01 | 2.0E-01 | 8.7E-01 | 7.7E-01 | 5.3E-02 | 9.2E-01 | 5.2E-01 |
| PE(P) 16:0/22:5 | 2.7E-01 | 2.8E-01 | 2.2E-01 | 6.9E-01 | 8.2E-01 | 3.3E-01 | 4.2E-01 | 3.0E-01 |
| PC C34:2 | 4.4E-01 | 6.0E-01 | 7.5E-01 | 6.5E-01 | 8.8E-01 | 4.3E-01 | 3.0E-01 | 3.1E-01 |
| PC C36:5 | 9.9E-01 | 3.2E-01 | 1.1E-01 | 8.9E-02 | 6.4E-02 | 4.4E-01 | 1.0E-01 | 1.0E-01 |
| PC C36:4 | 3.0E-01 | 6.5E-01 | 9.1E-01 | 9.1E-01 | 2.9E-01 | 5.4E-01 | 1.6E-01 | 2.2E-01 |
| PC C36:2 | 3.6E-01 | 8.9E-01 | 3.1E-01 | 9.6E-01 | 8.2E-01 | 1.9E-02 | 6.6E-01 | 9.2E-01 |
| PC C36:1 | 2.3E-01 | 9.7E-01 | 2.8E-01 | 8.8E-01 | 8.1E-01 | 9.4E-02 | 4.7E-01 | 2.0E-01 |
| PC C38:7 | 4.0E-01 | 2.8E-01 | 4.7E-02 | 2.1E-02 | 4.9E-02 | 1.2E-01 | 6.3E-02 | 1.4E-02 |
| PC C38:6 | 8.0E-01 | 6.7E-01 | 1.4E-01 | 1.5E-01 | 8.8E-01 | 3.7E-01 | 2.4E-01 | 2.6E-01 |
| PC C38:5 | 6.4E-01 | 7.3E-01 | 9.0E-01 | 6.0E-01 | 6.0E-01 | 8.3E-01 | 9.5E-02 | 2.2E-01 |
| PC C38:4 | 2.5E-01 | 2.2E-01 | 7.3E-01 | 6.3E-01 | 2.9E-01 | 9.6E-01 | 1.9E-01 | 1.9E-01 |
| PC C40:8 | 9.2E-01 | 9.9E-01 | 9.1E-01 | 4.3E-01 | 3.8E-01 | 5.6E-01 | 1.9E-01 | 3.1E-01 |
| PC C40:7 | 9.6E-01 | 8.8E-01 | 2.6E-01 | 1.5E-01 | 4.2E-01 | 7.5E-01 | 1.6E-01 | 2.9E-01 |
| PC C40:6 | 6.7E-01 | 8.5E-01 | 2.8E-01 | 1.0E-01 | 9.9E-01 | 6.8E-01 | 3.5E-01 | 2.6E-01 |
| PC C40:5 | 5.0E-01 | 4.8E-01 | 7.6E-01 | 7.4E-01 | 2.2E-01 | 7.6E-01 | 3.4E-01 | 3.3E-01 |
| PC C40:4 | 1.4E-01 | 7.4E-01 | 3.0E-01 | 5.4E-01 | 1.5E-01 | 5.7E-01 | 7.8E-01 | 3.6E-01 |

**Supplementary Results Table S2.** **Linear mixed effects model of lipids and the class (PD or control) and mitochondrial activity (complex I, II-III and IV ratioed to citrate synthase) interaction.** *The significant interactions post Benjamini-Hochberg multiple testing correction at FDR = 10% are reported. The lipids are represented by the 95% confidence interval, standard error and the non-adjusted p-value. No significant interactions were observed in complex IV.*

|  | **Complex I/CS** | **95% CI +/- SE** | **Raw p-value** |
| --- | --- | --- | --- |
| Complex I/citrate synthase ratio | Cer d18:1/20:1 | [-23.6, -5.2 +/- 4.7] | 4.7E-03 |
|  | Cer d18:1/22:1 | [-32.8, -8.4 +/- 6.2] | 2.7E-03 |
|  | Cer OH d18:1/24:0 | [-23.9, -3.4 +/- 5.2] | 1.3E-02 |
|  | HexCer d18:1/16:0 | [-20.9, -4.3 +/- 4.2] | 5.8E-03 |
|  | HexCer d18:1/18:1 | [-23.9, -5.3 +/- 4.7] | 4.5E-03 |
|  | HexSph | [-42.8, -13.3 +/- 7.5] | 8.6E-04 |
|  | Lyso-PC C16:0 | [5.6, 21.3 +/- 4.0] | 2.1E-03 |
|  | Lyso-PC C18:2 | [7.6, 31.0 +/- 6.0] | 3.5E-03 |
|  | Lyso-PC C20:4 | [5.4, 25.5 +/- 5.1] | 7.6E-03 |
|  | Lyso-PC C22:6 | [6.4, 27.1 +/- 5.3] | 1.1E-02 |
|  | Lyso-PE(P) C18:1 | [-35.3, -10.6 +/- 6.3] | 1.1E-03 |
|  | PE(P) 16:0/18:1 | [-39.2, -13.4 +/- 6.6] | 5.5E-04 |
|  | PE(P) 16:0/22:4 | [-35.3, -5.5 +/- 7.6] | 1.3E-02 |
|  | PE(P) 18:0/18:1 | [-35.3, -9.6 +/- 6.6] | 2.2E-03 |
|  | PE(P) 18:1/18:1 | [-39.8, -11.8 +/- 7.1] | 1.2E-03 |
|  | PE(P) 18:1/20:4 | [-40.7, -10.9 +/- 7.6] | 2.1E-03 |
|  | PE(P) 18:1/22:4 | [-34.8, -7.3 +/- 7.0] | 5.6E-03 |
|  | PE(P) 18:1/22:5 | [-32.3, -7.4 +/- 6.3] | 3.8E-03 |
| Complex II-III/citrate synthase ratio |  |  |  |
|  | Cer d18:1/22:6 | [-14.2, -4.6 +/- 2.4] | 4.7E-04 |

**Supplementary Methods**

**Supplementary Methods Table S1**. **Demographics of samples from controls and Parkinson’s disease in eight different brain regions**. *Numbers of each group, mean age and standard deviation, percentage of females, and the results from FDR-adjusted (5%) Student’s two-tailed t-test comparing the ages between the groups are presented. The PD samples further consisted of individuals with Braak stage classified as 3, 4, 5 or 6 – divided into “Braak 3 - 4” and “Braak 5 - 6” in the table. In the cases where the number of age observations are only one, the actual age is given in parentheses.*

| **Region** | **Class** | **n** | **Average age +/- standard deviation** | **% Females** | **p-value: Age: Total PD vs control** | **p-value: Age: Braak 3-4 vs control** |
| --- | --- | --- | --- | --- | --- | --- |
| Caudate | Control | 13 | 74.3 ± 8.7 | 54% | NS | NS |
|  | Total PD | 5 | 76.2 ± 8.5 | 60% |  |  |
|  | Braak 3 - 4 | 4 | 74.8 ± 9.1 | 50% |  |  |
|  | Braak 5 - 6 | 1 | (82.0) | 100% |  |  |
| Cerebellum | Control | 14 | 72.0 ± 7.8 | 36% | * | NS |
|  | Total PD | 16 | 78.6 ± 5.9 | 44% |  |  |
|  | Braak 3 - 4 | 10 | 78.6 ± 7.0 | 60% |  |  |
|  | Braak 5 - 6 | 6 | 78.5 ± 4.0 | 17% |  |  |
| Cingulate cortex | Control | 13 | 72.6 ± 7.7 | 46% | NS | NS |
|  | Total PD | 11 | 78.9 ± 6.7 | 64% |  |  |
|  | Braak 3 - 4 | 10 | 78.6 ± 7.0 | 60% |  |  |
|  | Braak 5 - 6 | 1 | (82.0) | 100% |  |  |
| Frontal cortex | Control | 12 | 70.7 ± 6.8 | 25% | * | NS |
|  | Total PD | 13 | 77.8 ± 6.1 | 38% |  |  |
|  | Braak 3 - 4 | 7 | 77.1 ± 7.7 | 57% |  |  |
|  | Braak 5 - 6 | 6 | 78.5 ± 4.0 | 17% |  |  |
| Parahippocampus | Control | 11 | 71.0 ± 6.8 | 36% | NS | NS |
|  | Total PD | 5 | 79.6 ± 7.7 | 60% |  |  |
|  | Braak 3 - 4 | 4 | 79.0 ± 8.8 | 50% |  |  |
|  | Braak 5 - 6 | 1 | (82.0) | 100% |  |  |
| Parietal cortex | Control | 15 | 72.4 ± 7.7 | 33% | NS | NS |
|  | Total PD | 7 | 78.4 ± 7.6 | 71% |  |  |
|  | Braak 3 - 4 | 6 | 77.8 ± 8.2 | 67% |  |  |
|  | Braak 5 - 6 | 1 | (82.0) | 100% |  |  |
| Putamen | Control | 13 | 71.6 ± 6.4 | 38% | * | NS |
|  | Total PD | 10 | 78.3 ± 6.8 | 40% |  |  |
|  | Braak 3 - 4 | 4 | 78.0 ± 10.5 | 75% |  |  |
|  | Braak 5 - 6 | 6 | 78.5 ± 4.0 | 17% |  |  |
| Temporal cortex | Control | 16 | 72.3 ± 7.3 | 38% | * | NS |
|  | Total PD | 11 | 78.9 ± 6.7 | 64% |  |  |
|  | Braak 3 - 4 | 10 | 78.6 ± 7.0 | 60% |  |  |
|  | Braak 5 - 6 | 1 | (82.0) | 100% |  |  |

**Supplementary Methods Table S2.** **Chromatographic separation parameters for the three analytical LC methods.** *The table shows the analytes, column chemistry and temperature, composition of mobile phase A and B, and the timings, flow rate and B percentage of the LC gradient.*

| **Analytes** | **Column chemistry** | **Mobile phase** | **LC method** |
| --- | --- | --- | --- |
| GM1, GM2, GM3, NANA, ceramide | BEH Amide, 1 x 150 mm, 1.7µm  Column temperature: 60 ̊C | A: 10 mM NH_4_COOH in 85% acetonitrile  B: 10 mM NH_4_COOH in 15% acetonitrile | Time [min] \| Rate[mL/min] \| %B; 0 \| 0.2 \| 0; 0.5 \| 0.2 \| 0; 8 \| 0.2 \| 30; 10 \| 0.2 \| 70; 10.5 \| 0.2 \| 80; 11.5 \| 0.2 \| 80; 11.6 \| 0.2 \| 0; 13.3 \| 0.2 \| 0; 15.83 \| 0.2 \| 0; 16 \| 0.2 \| 0 |
| HexCer, Hex2Cer, Gb3Cer, Gb4Cer, HexSph, Hex2Sph, Gb3Sph | BEH C8, 2.1 x 50 mm, 1.7 µm  Column temperature: 50 ̊C | A: 0.1% formic acid in water  B: 0.1% formic acid in methanol | Time [min] \| Rate[mL/min] \| %B; 0 \| 0.5 \| 50; 0.2 \| 0.5 \| 50; 2 \| 0.5 \| 100; 3 \| 0.5 \| 100; 3.1 \| 0.5 \| 50; 5 \| 0.5 \| 50 |
| PE, PC, PE(P), SM, lyso-PE, lyso-PC, lyso-PE(P), lyso-SM | BEH HILIC, 2.1 x 150 mm, 1.7 µm  Column temperature: ambient | A: 10 mM NH_4_CH_3_COOH in 95% acetonitrile,  B: 10 mM NH_4_CH_3_COOH in 50% acetonitrile | Time [min] \| Rate[mL/min] \| %B \|; 0 \| 0.5 \| 0 \|; 0.5 \| 0.5 \| 0 \|; 3.75 \| 0.5 \| 20 \|; 3.76 \| 0.5 \| 100 \|; 5.49 \| 0.5 \| 100 \|; 5.5 \| 0.8 \| 100 \|; 5.75 \| 0.8 \| 100 \|; 5.76 \| 0.8 \| 0 \|; 7.5 \| 0.8 \| 0 \|; 7.51 \| 0.5 \| 0 \|; 7.75 \| 0.5 \| 0 \| |

**Supplementary Methods Table S3.** **Multiple reaction monitoring transitions and collision energies.** *The table shows the transition from precursor to product ion, the cone and collision energies applied, and finally if detection was performed in positive (ESI+) or negative (ESI-) electrospray ionisation mode.*

| **Compound** | **Transition** | **Cone** | **Collision** | **Detection mode** |
| --- | --- | --- | --- | --- |
| Cer d18:1/16:0 | 536.5 > 280.26 | 100 | 30 | ESI- |
| Cer d18:1/16:1 | 534.49 > 278.25 | 100 | 30 | ESI- |
| Cer d18:1/18:0 | 564.54 > 308.3 | 100 | 30 | ESI- |
| Cer d18:1/18:1 | 562.52 > 306.28 | 100 | 30 | ESI- |
| Cer d18:1/20:0 | 592.57 > 336.33 | 100 | 30 | ESI- |
| Cer d18:1/20:1 | 590.55 > 334.31 | 100 | 30 | ESI- |
| Cer d18:1/20:4 | 584.5 > 328.26 | 100 | 30 | ESI- |
| Cer d18:1/20:5 | 582.49 > 326.25 | 100 | 30 | ESI- |
| Cer d18:1/22:0 | 620.6 > 364.36 | 100 | 30 | ESI- |
| Cer d18:1/22:1 | 618.58 > 362.34 | 100 | 30 | ESI- |
| Cer d18:1/22:5 | 610.52 > 354.28 | 100 | 30 | ESI- |
| Cer d18:1/22:6 | 608.5 > 352.26 | 100 | 30 | ESI- |
| Cer d18:1/24:0 | 648.63 > 392.39 | 100 | 30 | ESI- |
| Cer d18:1/24:1 | 646.61 > 390.37 | 100 | 30 | ESI- |
| Cer d18:1/24:2 | 644.6 > 388.36 | 100 | 30 | ESI- |
| Cer d18:1/24:6 | 636.54 > 380.3 | 100 | 30 | ESI- |
| Cer d18:1/26:5 | 666.58 > 410.34 | 100 | 30 | ESI- |
| Cer d18:1/26:6 | 664.57 > 408.33 | 100 | 30 | ESI- |
| Cer-OH d18:1/16:0 | 552.5 > 296.26 | 100 | 30 | ESI- |
| Cer-OH d18:1/18:0 | 580.53 > 324.29 | 100 | 30 | ESI- |
| Cer-OH d18:1/22:1 | 634.58 > 378.34 | 100 | 30 | ESI- |
| Cer-OH d18:1/24:0 | 664.62 > 408.38 | 100 | 30 | ESI- |
| Cer-OH d18:1/24:1 | 662.61 > 406.37 | 100 | 30 | ESI- |
| GM1 d18:1/16:0 | 1516.84 > 290.09 | 70 | 58 | ESI- |
| GM1 d18:1/18:0 | 1544.87 > 290.09 | 70 | 58 | ESI- |
| GM1 d18:1/18:1 | 1542.85 > 290.09 | 70 | 58 | ESI- |
| GM1 d18:1/20:0 | 1572.9 > 290.09 | 70 | 58 | ESI- |
| GM1 d18:1/20:1 | 1570.88 > 290.09 | 70 | 58 | ESI- |
| GM1 d18:1/22:0 | 1600.93 > 290.09 | 70 | 58 | ESI- |
| GM1 d18:1/24:1 | 1626.95 > 290.09 | 70 | 58 | ESI- |
| GM2 d18:1/16:0 | 1354.78 > 290.09 | 70 | 58 | ESI- |
| GM2 d18:1/18:0 | 1382.82 > 290.09 | 70 | 58 | ESI- |
| GM2 d18:1/18:1 | 1380.8 > 290.09 | 70 | 58 | ESI- |
| GM2 d18:1/20:0 | 1410.85 > 290.09 | 70 | 58 | ESI- |
| GM3 d18:1/18:0 | 1179.74 > 290.09 | 70 | 58 | ESI- |
| Gb3Sph | 786.48 > 282.36 | 72 | 32 | ESI+ |
| Hex2Sph | 624.4 > 282.36 | 35 | 16 | ESI+ |
| HexSph -28 | 434.37 > 236.32 | 92 | 16 | ESI+ |
| HexSph | 462.46 > 264.3 | 92 | 16 | ESI+ |
| Lyso-PC C16:0 | 496.34 > 184.07 | 30 | 20 | ESI+ |
| Lyso-PC C18:2 | 520.34 > 184.07 | 30 | 20 | ESI+ |
| Lyso-PC C18:1 | 522.36 > 184.07 | 30 | 20 | ESI+ |
| Lyso-PC C18:0 | 524.37 > 184.07 | 30 | 20 | ESI+ |
| Lyso-PC C20:4 | 544.34 > 184.07 | 30 | 20 | ESI+ |
| Lyso-PC C22:6 | 568.34 > 184.07 | 30 | 20 | ESI+ |
| Lyso-PC C22:5 | 570.36 > 184.07 | 30 | 20 | ESI+ |
| Lyso-PC C22:4 | 572.37 > 184.07 | 30 | 20 | ESI+ |
| Lyso-PE C18:2 | 478.29 > 337.27 | 30 | 20 | ESI+ |
| Lyso-PE C18:1 | 480.31 > 339.29 | 30 | 20 | ESI+ |
| Lyso-PE C20:4 | 502.29 > 361.27 | 30 | 20 | ESI+ |
| Lyso-PE C22:6 | 526.29 > 385.27 | 30 | 20 | ESI+ |
| Lyso-PE C22:5 | 528.31 > 387.29 | 30 | 20 | ESI+ |
| Lyso-PE C22:4 | 530.33 > 389.31 | 30 | 20 | ESI+ |
| Lyso-PE(P) C16:0 | 438.32 > 266.46 | 52 | 24 | ESI+ |
| Lyso-PE(P) C18:1 | 464.34 > 292.47 | 42 | 24 | ESI+ |
| Lyso-PE(P) C18:0 | 466.35 > 294.49 | 40 | 24 | ESI+ |
| Lyso-PE C16:0 | 454.29 > 313.27 | 30 | 20 | ESI+ |
| Lyso-SM C16:1 | 437.31 > 184.07 | 30 | 20 | ESI+ |
| Lyso-SM C18:1 | 465.35 > 184.07 | 30 | 20 | ESI+ |
| PPPCS (Lyso-SM 509) | 509.31 > 184.07 | 30 | 20 | ESI+ |
| NANA | 307.95 > 86.81 | 60 | 12 | ESI- |
| Gb3Cer d18:1/16:0 | 1046.63 > 884.66 | 124 | 64 | ESI+ |
| Gb3Cer d18:1/18:0 | 1074.68 > 912.71 | 124 | 64 | ESI+ |
| Gb4Cer d18:1/16:0 | 1249.74 > 884.68 | 125 | 52 | ESI+ |
| Hex2Cer d18:1/16:0 | 884.61 > 722.56 | 124 | 57 | ESI+ |
| Hex2Cer d18:1/18:0 | 912.64 > 750.59 | 124 | 57 | ESI+ |
| Hex2Cer d18:1/24:1 | 994.72 > 832.66 | 124 | 57 | ESI+ |
| Hex2Cer d18:1/24:0 | 996.73 > 834.68 | 124 | 57 | ESI+ |
| Hex2Cer-OH d18:1/24:2 | 1008.7 > 846.64 | 124 | 57 | ESI+ |
| Hex2Cer-OH d18:1/24:1 | 1010.71 > 848.66 | 124 | 57 | ESI+ |
| HexCer d18:1/16:1 | 720.7 > 558.7 | 125 | 52 | ESI+ |
| HexCer d18:1/16:0 | 722.67 > 560.64 | 125 | 52 | ESI+ |
| HexCer d18:1/18:1 | 748.6 > 586.6 | 125 | 52 | ESI+ |
| HexCer d18:1/18:0 | 750.6 > 588.6 | 125 | 52 | ESI+ |
| HexCer-OH d18:1/18:0 | 766.6 > 264.3 | 140 | 44 | ESI+ |
| HexCer-OH d18:1/20:0 | 794.6 > 264.3 | 140 | 44 | ESI+ |
| HexCer-OH d18:1/22:0 | 820.7 > 658.8 | 140 | 44 | ESI+ |
| HexCer d18:1/24:1 | 832.7 > 670.7 | 125 | 52 | ESI+ |
| HexCer d18:1/24:0 | 834.7 > 672.7 | 125 | 52 | ESI+ |
| HexCer-OH d18:1/24:2 | 846.7 > 684.8 | 140 | 44 | ESI+ |
| HexCer-OH d18:1/24:1 | 848.7 > 686.8 | 140 | 44 | ESI+ |
| HexCer-OH d18:1/24:0 | 850.7 > 688.8 | 140 | 44 | ESI+ |
| HexCer d18:1/26:1 | 860.73 > 698.76 | 125 | 52 | ESI+ |
| HexCer d18:1/26:0 | 862.7 > 700.8 | 125 | 52 | ESI+ |
| HexCer-OH d18:1/26:1 | 876.72 > 714.75 | 140 | 44 | ESI+ |
| HexCer-OH d18:1/26:0 | 878.74 > 716.77 | 140 | 44 | ESI+ |
| PC C34:2 | 758.57 > 184.07 | 30 | 20 | ESI+ |
| PC C36:5 | 780.55 > 184.07 | 30 | 20 | ESI+ |
| PC C36:4 | 782.57 > 184.07 | 30 | 20 | ESI+ |
| PC C36:2 | 786.6 > 184.07 | 30 | 20 | ESI+ |
| PC C36:1 | 788.62 > 184.07 | 30 | 20 | ESI+ |
| PC C38:7 | 804.55 > 184.07 | 30 | 20 | ESI+ |
| PC C38:6 | 806.57 > 184.07 | 30 | 20 | ESI+ |
| PC C38:5 | 808.59 > 184.07 | 30 | 20 | ESI+ |
| PC C38:4 | 810.6 > 184.07 | 30 | 20 | ESI+ |
| PC C40:8 | 830.57 > 184.07 | 30 | 20 | ESI+ |
| PC C40:7 | 832.59 > 184.07 | 30 | 20 | ESI+ |
| PC C40:6 | 834.6 > 184.07 | 30 | 20 | ESI+ |
| PC C40:5 | 836.62 > 184.07 | 30 | 20 | ESI+ |
| PC C40:4 | 838.63 > 184.07 | 30 | 20 | ESI+ |
| PE C34:2 | 716.52 > 575.5 | 30 | 20 | ESI+ |
| PE C34:1 | 718.54 > 577.52 | 30 | 20 | ESI+ |
| PE C36:5 | 738.51 > 597.49 | 30 | 20 | ESI+ |
| PE C36:4 | 740.52 > 599.5 | 30 | 20 | ESI+ |
| PE C36:3 | 742.54 > 601.52 | 30 | 20 | ESI+ |
| PE C36:2 | 744.55 > 603.54 | 30 | 20 | ESI+ |
| PE C36:1 | 746.57 > 605.55 | 30 | 20 | ESI+ |
| PE C38:7 | 762.51 > 621.49 | 30 | 20 | ESI+ |
| PE C38:6 | 764.52 > 623.5 | 30 | 20 | ESI+ |
| PE C38:5 | 766.54 > 625.52 | 30 | 20 | ESI+ |
| PE C38:4 | 768.55 > 627.54 | 30 | 20 | ESI+ |
| PE C40:8 | 788.52 > 647.5 | 30 | 20 | ESI+ |
| PE C40:7 | 790.54 > 649.52 | 30 | 20 | ESI+ |
| PE C40:6 | 792.55 > 651.54 | 30 | 20 | ESI+ |
| PE C40:5 | 794.57 > 653.55 | 30 | 20 | ESI+ |
| PE C40:4 | 796.59 > 655.57 | 30 | 20 | ESI+ |
| PE(P) 18:0/20:4 | 752.56 > 361.27 | 76 | 24 | ESI+ |
| PE(P) 16:0/22:4 | 752.56 > 389.31 | 76 | 24 | ESI+ |
| PE(P) 18:1/22:6 | 774.54 > 385.27 | 76 | 24 | ESI+ |
| PE(P) 18:0/22:6 | 776.56 > 385.27 | 76 | 24 | ESI+ |
| PE(P) 18:1/22:5 | 776.56 > 387.29 | 76 | 24 | ESI+ |
| PE(P) 18:0/22:5 | 778.58 > 387.29 | 76 | 24 | ESI+ |
| PE(P) 18:1/22:4 | 778.58 > 389.31 | 76 | 24 | ESI+ |
| PE(P) 18:0/22:4 | 780.59 > 389.31 | 76 | 24 | ESI+ |
| PE(P) 18:1/16:0 | 702.54 > 313.27 | 76 | 24 | ESI+ |
| PE(P) 16:0/18:1 | 702.54 > 339.29 | 76 | 24 | ESI+ |
| PE(P) 16:0/20:4 | 724.53 > 361.27 | 76 | 24 | ESI+ |
| PE(P) 18:1/18:1 | 728.56 > 339.29 | 76 | 24 | ESI+ |
| PE(P) 18:0/18:1 | 730.58 > 339.29 | 76 | 24 | ESI+ |
| PE(P) 16:0/22:6 | 748.53 > 385.27 | 76 | 24 | ESI+ |
| PE(P) 18:1/20:4 | 750.54 > 361.27 | 76 | 24 | ESI+ |
| PE(P) 16:0/22:5 | 750.54 > 387.29 | 76 | 24 | ESI+ |
| SM C34:2 | 701.58 > 184.07 | 30 | 20 | ESI+ |
| SM C34:3 | 699.58 > 184.07 | 30 | 20 | ESI+ |
| SM C36:3 | 727.61 > 184.07 | 30 | 20 | ESI+ |
